# Localized heme sensing through a ternary molecular glue

**DOI:** 10.64898/2026.05.07.723605

**Authors:** Michael Heider, Clara Hipp, Zhi Yang, Han Xiao, Tobias Beschauner, Eddie Wehri, Wencke Walter, Rumi Sherriff, Srividya Chandrasekhar, Torsten Haferlach, Julia Schaletzky, Michael Rapé

**Affiliations:** Department of Molecular and Cell Biology, University of California at Berkeley, Berkeley, CA 94720, USA; Howard Hughes Medical Institute, University of California at Berkeley, Berkeley, CA 94720, USA; Molecular Therapeutics Initiative, University of California at Berkeley, Berkeley, CA 94720, USA; MLL Munich Leukemia Laboratory, Munich, Germany; California Institute for Quantitative Biosciences (QB3), University of California at Berkeley, Berkeley, CA 94720, USA; Present address: Research Institute of Molecular Pathology (IMP), Vienna BioCenter (VBC), Vienna, Austria

**Author notes:** These authors contributed equally (Clara Hipp, Zhi Yang).

**Keywords:** ubiquitin, molecular glue, FEM1B, mitochondria, heme, AML

## Abstract

Molecular glues are an emerging class of therapeutics that stabilize binary interactions and thereby rewire disease-relevant protein networks. Whether glues can integrate additional information to orchestrate signaling beyond initial complex formation is unknown. Here, we report that cells use an endogenous glue strategy to sense heme, an essential metabolite with deleterious pro-oxidant properties. Distinct from other glues, heme bridges three polypeptides to trigger degradation of the transcriptional repressor BACH1 through cytoplasmic, but not mitochondrial, CUL2^FEM1B^. This mechanism allows cells to eliminate toxic heme in the cytoplasm by inducing expression of the heme-degrading oxygenase HMOX1, yet ignore mitochondrial heme destined for function in the electron transport chain. While protective in healthy cells, ternary glue signaling creates a therapeutic vulnerability for Acute Myeloid Leukemias dependent on high rates of ETC assembly. Molecular glues can therefore drive assembly of higher-order complexes to establish localized signaling, which offers unexplored opportunities for induced proximity therapeutics.

## Introduction

Molecular glues are an emerging class of therapeutics that promote specific interactions to rewire disease-relevant signaling pathways ^1–5^. As glues bind at the interface between proteins, they do not require preexisting drug pockets and can therefore target factors previously deemed undruggable. The first glues, rapamycin and FK506, were found to drive the association of chaperones with mTORC1 kinase and calcineurin phosphatase, respectively, thereby modulating phosphorylation networks at the heart of immunosuppression ^6,7^. It was later realized that compounds, such as thalidomide or indisulam, recruit E3 ligases to induce the ubiquitylation and proteasomal degradation of gene expression regulators ^8–16^. Phenotypic screens have since identified a wide range of additional effectors ^1,5^, yet whether molecular glues only differ in the proteins they recruit or may have properties beyond complex formation is still not known.

Akin to therapeutics, endogenous molecules can stabilize specific interactions and thus act as glues. The plant hormones auxin, jasmonate, gibberellic acid and strigolactone tether transcriptional repressors to E3 ligases, unleashing gene expression programs that govern organismal growth, root formation, or flowering ^17–20^. In human cells, purine monophosphates were recently shown to glue the rate-limiting enzyme of purine synthesis, PPAT, to its inhibitor NUDT5 ^21^. The discovery of this metabolite glue signaling revealed how cells adjust purine levels to fluctuating demands, but it also informed on the mechanism of chemotherapeutics that were introduced into the clinic ∼75 years ago. Mirroring endogenous glues, thioguanine and mercaptopurine drugs were found to stabilize the PPAT-NUDT5 complex and exert their effects by starving cancer cells of purines required for DNA replication, energy metabolism, or signal transduction ^21–24^. Endogenous molecular glues therefore regulate crucial signaling networks that can be modulated for therapeutic benefit. Unfortunately, only very few endogenous glues are known, and pivotal mechanisms of glue signaling likely remain to be discovered.

Given their small size, molecular glues typically contribute only limited binding energy to complex formation and predominantly act by stabilizing weak preexisting interactions ^2,15^. Glue-induced complex formation accordingly depends on complementary binding surfaces and occurs through cooperative interactions between all partners, which together establishes the high specificity of this signaling modality ^7,17,25–27^. It is widely thought that molecular glues integrate these features by bridging interactions between two proteins in a constitutive manner. Whether glues can incorporate additional information about the cellular state is unknown, and the full potential of molecular glues for endogenous signaling or therapeutic application therefore remains to be established.

Here, we report that cells use an endogenous glue strategy to monitor heme, an essential co-factor of the electron transport chain (ETC) that when present in excess exerts severe oxidative stress. Different from known glues, heme bridges three polypeptides to trigger the selective degradation of the transcriptional repressor BACH1 through cytoplasmic, but not mitochondrial, CUL2^FEM1B^. Heme is, therefore, a ternary molecular glue. Cells use this mechanism to counteract toxic accumulation of heme in the cytoplasm, while ignoring mitochondrial heme required for ETC assembly and function. Reflecting its protective function, disrupting ternary glue signaling creates therapeutic opportunities in Acute Myeloid Leukemias that rely on increased assembly of the heme-dependent ETC. Our work reveals localized signaling by a small molecule, and it illustrates how identifying endogenous molecular glues can rapidly inform therapeutic strategies against diseases of high unmet need.

## Results

### The stress response E3 ligase CUL2^FEM1B^ is functionally linked to heme metabolism

We initiated this study based on a need for new therapeutic approaches for Acute Myeloid Leukemia (AML), a cancer that remains difficult to treat in elderly patients and those with relapsed or refractory disease ^28^. Despite genetic heterogenicity, aggressive AML subtypes share a requirement for oxidative phosphorylation driven by mitochondrial electron transport chain (ETC) complexes ^29–32^. This observation motivated development of inhibitors against ETC complex I, which showed promising preclinical data yet failed due to toxicity and resulting inadequate dosing ^33,34^. We speculated that targeting regulators of the ETC that are particularly important for AML, rather than core ETC subunits required by all cells, may provide more effective therapeutic opportunities.

We therefore assessed the role of the E3 ligase CUL2^FEM1B^ in AML. CUL2^FEM1B^ is activated when ETC levels drop below cellular needs, triggering protein degradation at TOM complexes to increase mitochondrial protein import and thereby restore assembly of the rate-limiting ETC complex IV (cIV) ^35–37^. We deleted the specificity factor of CUL2^FEM1B^, its substrate adaptor FEM1B, in cell lines derived from mitochondrially active AML subtypes ^31^ and monitored cell fitness using competition approaches ^38,39^. In line with a crucial role of CUL2^FEM1B^, AML cells lacking *FEM1B* were efficiently depleted from mixtures with their WT counterparts (**Figure 1A; Figure S1A**). To provide mechanistic insight into the underlying regulation, we conducted whole genome synthetic lethality screens to compare effects of gene deletions onto WT and *ΔFEM1B* AML cells (**Figure 1B**). As expected, mitochondrial processes were essential for WT AML cells (**Figure S1B**), and deletion of genes encoding ETC subunits, ETC assembly factors, or mitochondrial translation factors had much stronger effects in *ΔFEM1B* than in WT cells (**Figure 1B**). We validated these synthetic lethal interactions by direct cell competition (**Figure 1C**). These results confirmed that CUL2^FEM1B^ is an important enzyme in AML that acts, at least in part, by modulating the ETC.

**Figure 1:**
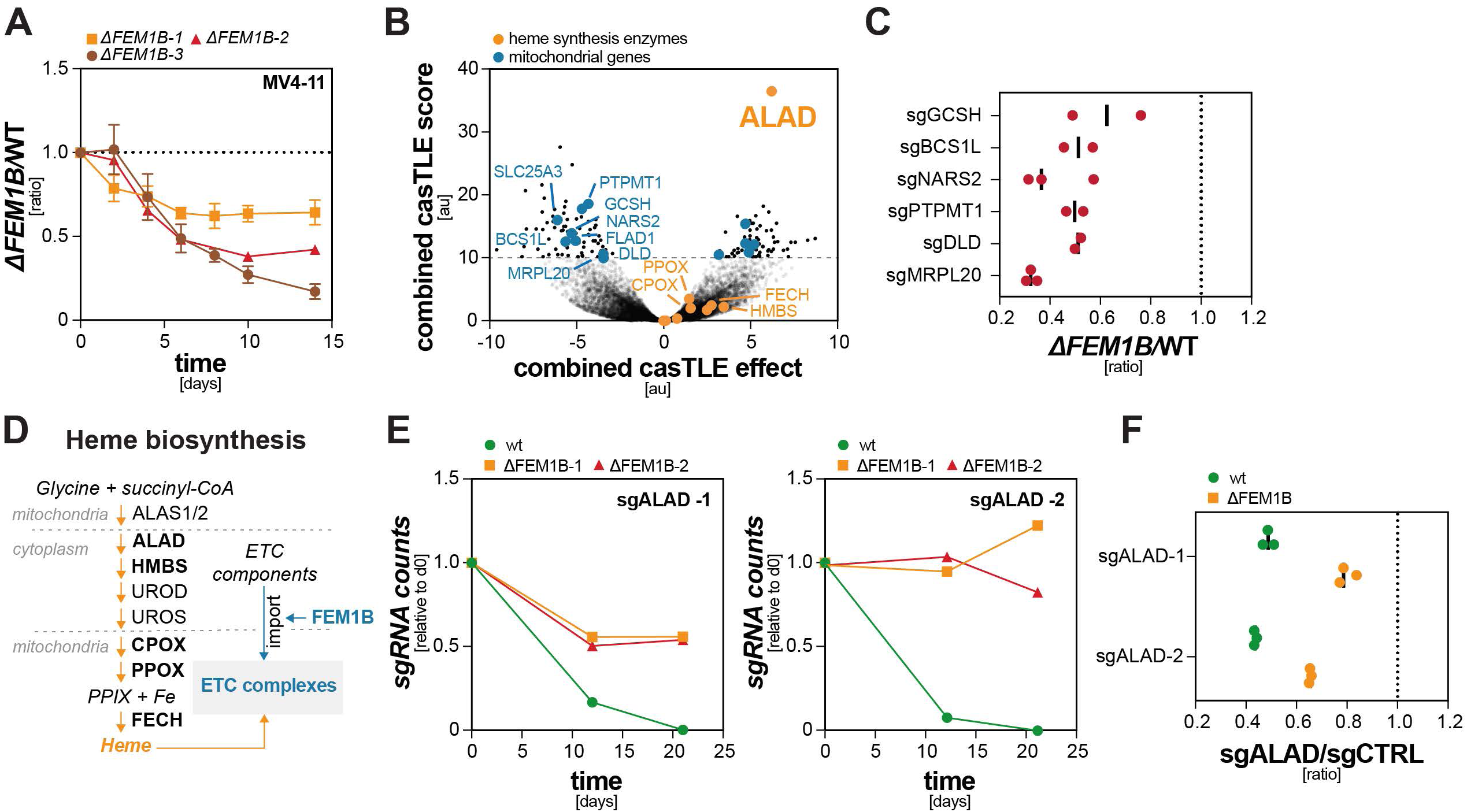
The E3 ligase CUL2^FEM1B^ is functionally linked to heme metabolism. **A.** FEM1B preserves AML cell fitness. Three mCherry-expressing *ΔFEM1B* MV4-11 clones were mixed with GFP-expressing wt MV4-11 cells and followed by flow cytometry for 14d. Datapoints represent mean ± S.E.M. of n=3 independent experiments. **B.** Whole genome dropout CRISPR screen reveals genetic interactors of FEM1B in AML cells. Shown is the CasTLE analysis ^86^ of change in sgRNA abundance over 21 days in two distinct *ΔFEM1B* MV4-11 clones compared to wt MV4-11 cells. Dashed line indicates 10% FDR. **C.** Loss of mitochondrial CRISPR screen hits is synthetic lethal in MV4-11 *ΔFEM1B* cells. GFP-expressing WT cells were mixed with mCherry-expressing *ΔFEM1B* cells, depleted of indicated mitochondrial genes, and ratios were determined after 12d of cell competition. Datapoints represent n=2-3 independent experiments. **D.** Schematic of the heme biosynthesis pathway. CUL2^FEM1B^ regulates mitochondrial import required for assembly of the heme-containing ETC complex IV ^36^. **E.** ALAD-dependency of AML cells is strongly attenuated in *ΔFEM1B* cells. Relative counts of two sgRNAs (normalized to d0) targeting the heme biosynthesis enzyme ALAD throughout the 21d CRISPR screen. **F.** Cell competition assays confirm increased tolerance of *ΔFEM1B* cells to ALAD loss. Fluorescently labelled sgALAD or sgCTRL cells were assessed for MV4-11 wt and MV4-11 *ΔFEM1B* cells, respectively, and followed by flow cytometry for 12d. Datapoints represent n=3 independent experiments.

In addition to these expected results, our screens revealed a surprising enrichment of sgRNAs targeting δ-amino-levulinate dehydratase (ALAD) in *ΔFEM1B* cells (**Figure 1B, D**). ALAD catalyzes the second committed condensation step in heme biosynthesis ^40^. sgRNAs against most other heme biosynthetic enzymes were also enriched in *ΔFEM1B* compared to WT cells, suggesting that inhibition of heme synthesis is tolerated much better when CUL2^FEM1B^ is inactive. Consistent with this notion, single guide analyses showed that deletion of *FEM1B* protected AML cells against toxicity caused by the loss of ALAD (**Figure 1E**), and independent competitions confirmed that *ΔFEM1B* cells tolerated ALAD depletion much better than WT cells (**Figure 1F**). This synthetic viability phenotype was specific for AML subtypes with high ETC activity, as it was not observed in a parallel screen conducted in cells that are less dependent on oxidative phosphorylation (**Figure S1C, D**). These observations suggested that CUL2^FEM1B^ exerts a second, hitherto unknown role in controlling heme metabolism that is related to its role in safeguarding ETC function.

Heme is an iron-containing porphyrin that acts as an essential co-factor for ETC complexes, including cIV ^41,42^. While lack of heme impairs mitochondrial function, excess heme is a strong oxidant that disrupts protein and membrane integrity ^43^; cells must therefore ensure that heme levels stay within an optimal range. Heme is modified for incorporation into cIV by the mitochondrial proteins COX10 and COX15 whose activity is coupled to cIV assembly intermediates ^44–46^. Delays in cIV formation result in excess free heme, which diffuses into the cytoplasm to elicit stress; cells must therefore also coordinate heme synthesis with cIV assembly. As CUL2^FEM1B^ appeared to impact both heme metabolism and cIV assembly, we speculated that modulating this E3 ligase could create therapeutic vulnerabilities in AML. To guide any treatment strategies, we decided to investigate how CUL2^FEM1B^ controls heme biology.

### CUL2^FEM1B^ induces heme-dependent degradation of BACH1

We started out by searching for ubiquitylation targets of CUL2^FEM1B^ with functions in heme metabolism. We therefore purified a substrate-trapping variant, FEM1B^R126A/L597A^, from two AML cell lines and determined binding partners by mass spectrometry. In addition to known targets, such as FNIP1 or COA4, these experiments identified the transcriptional repressor BACH1 as a specific binder of FEM1B (**Figure 2A; Figure S2A**). Immunoprecipitation followed by Western blotting confirmed the interaction of FEM1B with BACH1 in a different cell type, indicating that it is not constrained to AML (**Figure 2B**). Reciprocal analysis of BACH1 immunoprecipitates validated its prominent association with FEM1B (**Figure 2C**). These studies also confirmed binding of BACH1 to two E3 ligases, CUL1^FBXO22^ and CUL1^FBXL17^, that mediate BACH1 degradation after it had been damaged by heme-induced oxidative stress ^47–50^.

**Figure 2:**
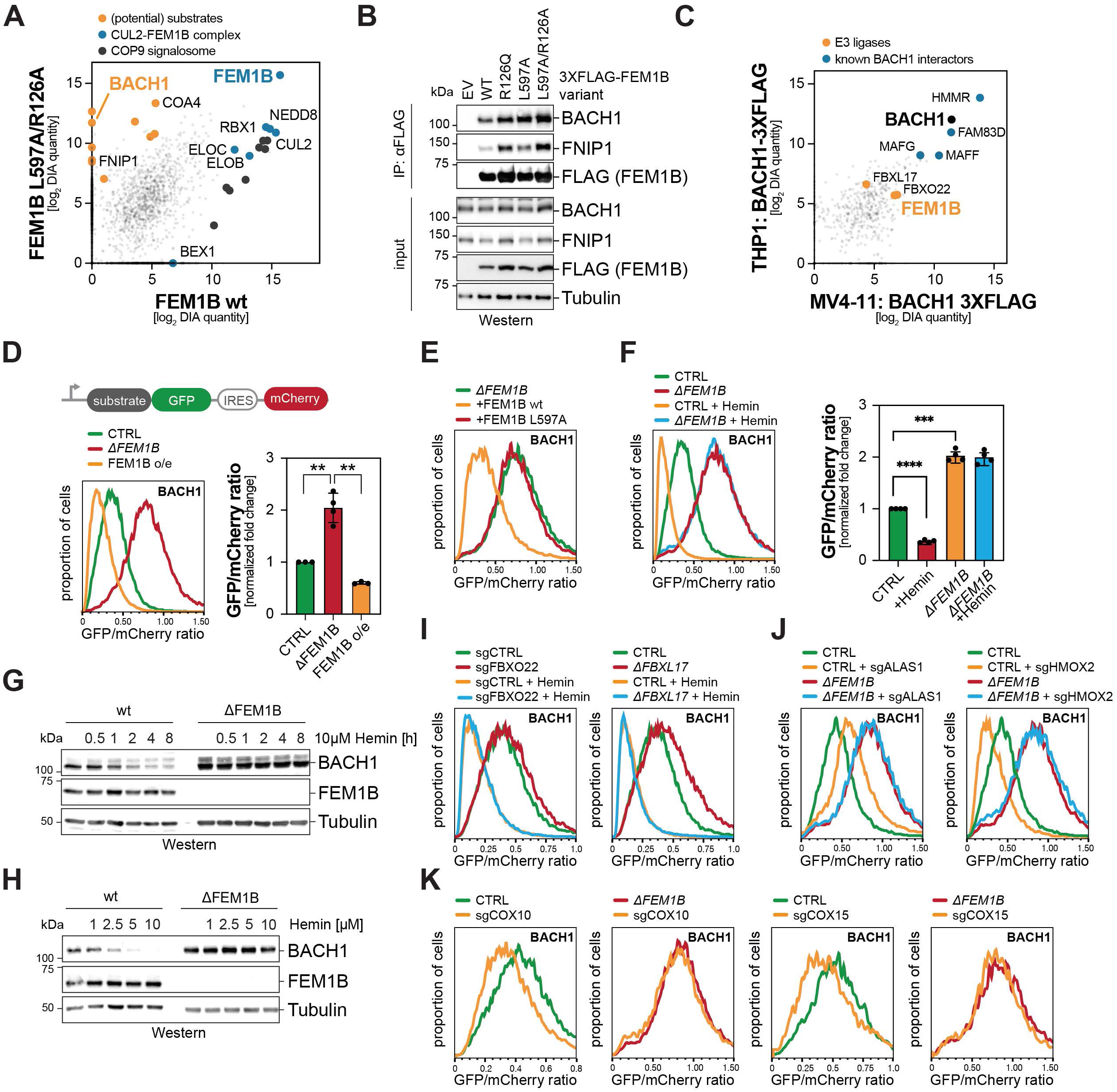
CUL2^FEM1B^ induces heme-dependent degradation of BACH1. **A.** Identification of BACH1 as a potential substrate of CUL2^FEM1B^ by immunoprecipitation of lentivirally expressed ^3XFLAG^FEM1B or substrate trap mutant ^3XFLAG^FEM1B^R126A/L597A^ from MV4-11 cells coupled to mass spectrometry. Candidate substrates are enriched in affinity-purifications of FEM1B^R126A/L597A^. **B.** Validation of the interaction between FEM1B and BACH1. FLAG-tagged FEM1B variants were immunoprecipitated from HEK293T cells and interactors were detected by Western blotting using the indicated antibodies. Similar results in n=2 independent experiments. **C.** Immunoprecipitation of BACH1^3XFLAG^ identifies FEM1B as a prominent interactor. BACH1^3XFLAG^ was lentivirally expressed in MV4-11 and THP1 cells and immunoprecipitated proteins were detected by mass spectrometry. **D.** BACH1 degradation requires FEM1B. BACH1 stability reporters were expressed in wt or *ΔFEM1B* HEK293T cells in the presence or absence of overexpressed FEM1B. *Upper scheme:* reporter construct with substrate (BACH1) fused to GFP co-expressed with mCherry under control of an internal ribosome entry site (IRES). Quantification of median fluorescence intensity ratios (MFI) from n=3-4 independent experiments is shown to the right. **E.** WT, but not the CUL2-binding deficient FEM1B^L597A^, restores BACH1 degradation in *ΔFEM1B* HEK293T cells. Similar results in n=3 independent experiments. **F.** BACH1 degradation upon Hemin treatment depends on FEM1B. BACH1 reporter constructs were transiently transfected into WT or *ΔFEM1B* HEK293T cells, and cells were treated with 10µM hemin for 16h. *Right:* Quantification of mean fluorescence intensity ratios from n=4 independent experiments. **G.** Heme-induced turnover of endogenous BACH1 is dependent on FEM1B in AML cells. WT and *ΔFEM1B* MV4-11 cells were treated 10µM hemin for the indicated times, and endogenous protein expression was assessed by Western blotting using the indicated antibodies. Similar results were obtained in n=3 independent experiments. **H.** WT and *ΔFEM1B* MV4-11 cells were treated with increasing concentrations of hemin for 16h, and protein levels were determined using Western blotting. Similar results in n=3 independent experiments. **I.** FBXO22 and FBXL17 do not mediate heme-induced degradation of BACH1. BACH1 stability reporters were transiently transfected into HEK293T that were depleted of either *FBXO22* or *FBXL17*, and cells were treated with 10µM hemin for 16h. Similar results in n=2 independent experiments. **J.** Manipulation of endogenous heme levels modulates BACH1 stability dependent on FEM1B. ALAS1 or HMOX2 were depleted from WT or *ΔFEM1B* HEK293T cells stably expressing a BACH1 stability reporter, and BACH1 levels were determined by flow cytometry. Similar results in n=3 independent experiments. **K.** Disruption of heme integration into ETC cIV destabilizes BACH1 dependent on FEM1B. COX10 or COX15 were depleted from WT or *ΔFEM1B* HEK293T cells stably expressing the BACH1 stability reporter, and BACH1 levels were determined by flow cytometry. Similar results in n=3 independent experiments. Statistical significance in (D) and (F) was determined using one-sample t-tests (** p<0.01, *** p<0.001 and **** p<0.0001).

Having identified BACH1 as an interactor of CUL2^FEM1B^, we used a flow cytometry strategy to assess whether it is degraded through this E3 ligase ^51,52^. We found that deletion of *FEM1B* strongly stabilized BACH1, while its overexpression had the opposite effect of lowering BACH1 levels (**Figure 2D**). Other newly identified binding partners of FEM1B in AML cells did not behave in this manner (**Figure S2B**). FEM1B^L597A^, which fails to bind CUL2, did not elicit BACH1 turnover (**Figure 2E**), and BACH1 was stabilized by dominant-negative CUL2, the Cullin-RING E3 ligase inhibitor MLN4924, or the proteasome inhibitor carfilzomib (**Figure S2C, D**). Further underscoring the specificity of this proteolytic circuit, FEM1B did not turn over the related BACH2 (**Figure S2E**), while the close homolog of FEM1B, FEM1A, was unable to degrade BACH1 (**Figure S2F**). Thus, CUL2^FEM1B^ not only binds BACH1, but also targets this transcriptional repressor for proteasomal degradation. While this manuscript was in preparation, an independent study also suggested that CUL2^FEM1B^ may target BACH1 for degradation ^53^.

Suggesting that it could mediate effects of CUL2^FEM1B^ onto heme metabolism, BACH1 had previously been shown to be degraded in response to heme accumulation ^54–56^. Recent work had identified two E3 ligases, SCF^FBXL17^ and SCF^FBXO22^, that degrade BACH1 molecules damaged by heme-dependent oxidative stress ^47–50^. To test if CUL2^FEM1B^ contributes to the turnover of BACH1 in response to heme, we supplemented cells with a membrane-permeable analog, hemin, and measured its stability by flow cytometry. While hemin induced degradation of BACH1, we were surprised to see that this was entirely dependent on *FEM1B* (**Figure 2F; Figure S2G**). Hemin also depleted endogenous BACH1 in AML cells in a time- and dose-dependent manner, which was abolished if cells lacked *FEM1B* (**Figure 2G, H**). Other CUL2^FEM1B^ substrates, including FNIP1 and COA4, were not affected by changes in heme levels (**Figure S2I**), and SCF^FBXO22^ and SCF^FBXL^^17^ did not drive heme-induced degradation under these conditions (**Figure 2I; Figure S2J-L**). CUL2^FEM1B^ is therefore the sole E3 ligase that promotes acute BACH1 turnover in response to rising heme levels.

As hemin does not reflect physiology, we next assessed if CUL2^FEM1B^ also controls BACH1 degradation upon changes in endogenous heme. We first lowered heme by depleting the biosynthetic enzymes ALAD or ALAS1 and found that such treatments stabilized BACH1 (**Figure 2J; Figure S2M**). We then increased heme by depleting the constitutively expressed heme-degrading oxygenase HMOX2 ^57^ or by overexpressing ALAD or ALAS1. These conditions accelerated BACH1 turnover, and this was fully dependent on FEM1B (**Figure 2J; Figure S2N**). Finally, we raised free heme levels by impairing its incorporation into nascent cIV by depleting COX10, COX15, or other assembly factors; this treatment also triggered BACH1 degradation dependent on continued heme synthesis (**Figure 2K; Figure S2O, P**). Cells therefore react to increased endogenous heme levels by triggering acute BACH1 degradation through CUL2^FEM1B^.

### Cryo-EM structure of CUL2^FEM1B^ bound to BACH1 and heme

By assessing truncation variants, we identified a carboxy-terminal region in BACH1, referred to as BACH1^CT^, that was necessary and sufficient for its heme-dependent interaction with FEM1B (**Figure 3A**). The same domain was necessary and sufficient for heme-dependent degradation of BACH1 through CUL2^FEM1B^ (**Figure 3B**). BACH1 contains six Cys/Pro motifs known to engage heme ^54^ (**Figure S3A**), and only inactivation of the Cys/Pro motif within BACH1^CT^ rendered BACH1 insensitive to FEM1B-binding and heme-induced degradation (**Figure 3C; Figure S3B**). Deletion or mutation of BACH1’s BTB domain, which is recognized by SCF^FBXO22^ and SCF^FBXL1747,48,58,59^, did not affect its heme-induced degradation (**Figure S3C**). Thus, CUL2^FEM1B^ recognizes a specific motif in the C-terminus of BACH1 to instigate its heme-dependent degradation.

**Figure 3:**
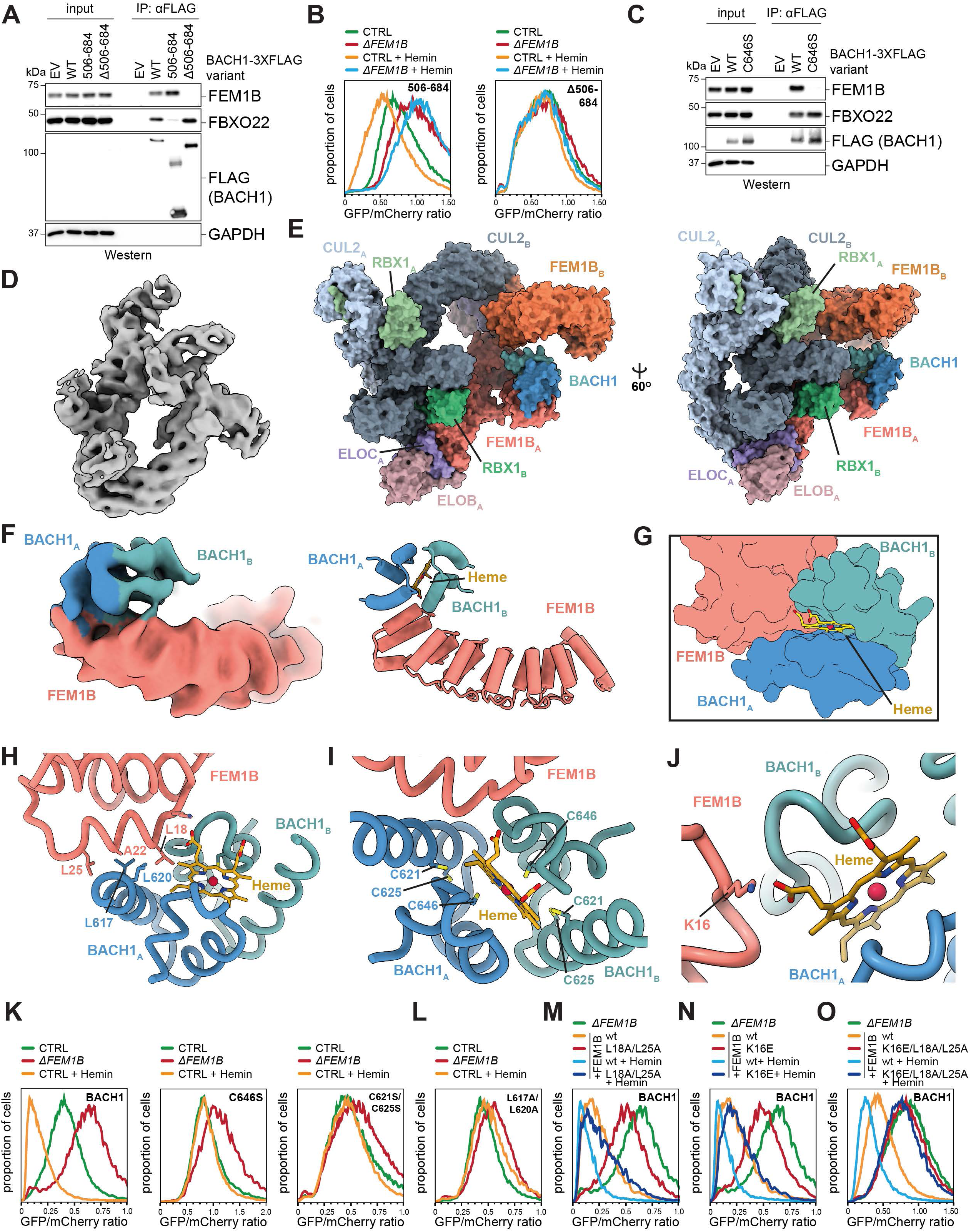
Cryo-EM structure of CUL2^FEM1B^ bound to BACH1 and heme. **A.** A carboxy-terminal domain in BACH1 (BACH1^CT^; residues 506-684) is required and sufficient for binding to FEM1B. BACH1^3XFLAG^ constructs were transiently transfected into *ΔBACH1* HEK293T cells and immuno-precipitates were analyzed for co-purifying proteins by Western blotting. Similar results in n=3 independent experiments. **B.** BACH1^CT^ is sufficient and required for heme-dependent degradation by FEM1B. BACH1 stability reporters were transiently transfected into WT and *ΔFEM1B* HEK293T cells, and cells were treated with 10µM hemin for 16h. Similar results in n=3 independent experiments. **C.** BACH1^C646^ is required for binding to FEM1B. BACH1^3XFLAG^ variants were transiently transfected into *ΔBACH1* HEK293T cells, and immunoprecipitates were analyzed by Western blotting. Similar results in n=3 independent experiments. **D.** BACH1-bound CUL2^FEM1B^ forms an asymmetric dimer. Cryo-EM density map of the dimeric CUL2-RBX1-ELOB/C-FEM1B-BACH1^CT^ complex, Electron Microscopy Data Bank (EMDB): EMD-76876, contour level 0.03) **E.** Surface representation of the complex reveals structural details of the dimeric assembly and highlights positioning of BACH1 in the active complex. Different shades of the same colors used for individual subunits in the two protomers. Subscripted A/B denotes the corresponding protomer. Right: 60° rotation to the right. **F.** BACH1 binds to the cap of the FEM1B N-terminal ankyrin repeats. *Left:* Cryo-EM density map of the locally refined FEM1B-BACH1 binding interface (PDB 12ZX/ EMD-76910, contour level 0.026). *Right:* AF3 model of the BACH1-FEM1B binding interface. **G.** The ternary molecular glue heme enables an expansive binding interface between two BACH1 subunits and amino-terminal ankyrin repeats of FEM1B by bridging three polypeptide chains. **H.** Hydrophobic interactions in both FEM1B and BACH1^CT^ mediate binding. AF3 model of the FEM1B-BACH1 interface, guided by cryo-EM data and highlighting hydrophobic residues L18, A22, L25 of FEM1B and L617, L620 of BACH1. **I.** Heme facilitates dimerization of two BACH1 protomers. Cryo-EM guided AF3 model of two BACH1 protomers sandwiching heme, highlighting the heme-coordinating residues C646, C621 and C625. **J.** BACH1-bound heme directly engages FEM1B. Cryo-EM guided AF3 model of the FEM1B-BACH1 interface highlighting K16 interaction with the carboxyl group of the porphyrin ring. **K.** Mutation of heme coordinating residues in BACH1 inhibits its heme-induced degradation via CUL2^FEM1B^. BACH1, BACH1^C646S^ or BACH1^C621S/C625S^ stability reporters were transiently transfected into WT or *ΔFEM1B* HEK293T cells, and cells were treated with 10µM hemin for 16h. BACH1 abundance was monitored by flow cytometry. Similar results in n=3 independent experiments. **L.** Mutation of hydrophobic BACH1 residues at the interface with FEM1B inhibits its heme-induced degradation. BACH1^L617A/L620A^ stability reporters were transiently transfected into WT or *ΔFEM1B* HEK293T cells, cells were treated with 10µM hemin for 16h, and BACH1 abundance was monitored by flow cytometry. Similar results in n=3 independent experiments. **M-O.** Mutation of FEM1B residues involved in BACH1 or heme binding disrupt BACH1 degradation. FEM1B variants and BACH1 stability reporters were transiently transfected into *ΔFEM1B* HEK293T cells, and cells were treated with 10µM hemin for 16h. Similar results in n=3 independent experiments.

Having found this degron, we reconstituted a complex composed of CUL2^FEM1B^, BACH1^CT^, and heme and subjected it to single-particle cryo-EM analysis (**Figure S4A, B; Figure S5A, B**). Despite significant flexibility in CUL2^FEM1B^, as reported ^60^, we could combine this data with Alpha-Fold3 to derive a structural model for the heme-dependent assembly (**Figures 3D, E**). When engaged to BACH1, CUL2^FEM1B^ forms an asymmetric dimer that is similar in organization to CUL2^FEM1B^ complexes bound to the candidate substrate PLD6 or the mitochondrial anchor TOM20^61^ (**Figures 3D, E**). Our structure highlighted that dimerization of CUL2^FEM1B^ requires previously identified residues in the TPR domain of FEM1B ^60^. Mutating these residues prevented the heme-induced degradation of BACH1 (**Figure S5C**), indicating that E3 ligase dimerization is important for turning over this substrate.

As the flexibility of CUL2^FEM1B^ limited the resolution of this structure, we repeated our analyses using carbon-film coated grids. While this strategy interfered with CUL2^FEM1B^ dimerization, it improved overall resolution to 3.8Å and, combined with a local refinement map, allowed us to assign BACH1 into the density map (**Figure 3F; Figure S5D, E**). Strikingly, this revealed that CUL2^FEM1B^ engaged BACH1 at a position different from all known substrates. While other targets of CUL2^FEM1B^ bind a conserved groove in FEM1B formed by its ankyrin repeats and TPR-domain ^35,37,60–62^, BACH1 associates with the same amino-terminal region of FEM1B that is recognized by its mitochondrial anchor TOM20 and that is required for targeting CUL2^FEM1B^ to outer mitochondrial membranes ^36^ (**Figure 3F, G; Figure S5F**). This structurally defined interaction surface for BACH1 differs from the binding site suggested by deletion studies, likely because the respective deletions in the latter work destabilize the FEM1B fold and disrupt E3 ligase dimerization ^53^. Thus, BACH1 accesses a site in FEM1B that is typically reserved for E3 ligase regulation, not substrate binding.

In addition to its unique binding site, BACH1 engages FEM1B in an oligomeric state that differs from known CUL2^FEM1B^ substrates. While canonical substrates bind using a single degron, BACH1 engages the E3 ligase as a dimer of BACH1^CT^ subunits that wrap around the amino-terminal cap of FEM1B (**Figure 3F, G**). One BACH1^CT^ subunit binds FEM1B through hydrophobic interactions between an α-helix centered on Leu residues in BACH1 (L617/L620) and Leu residues in the amino-terminal helix of FEM1B (L18/L25) (**Figure 3H**). As discussed below, the same FEM1B leucines are required for recognition of TOM20 ^36^. The second subunit of the BACH^CT^ dimer docks against the backside of the FEM1B ankyrin repeats, an interaction that is mediated by FEM1B-K60 and BACH1-Y641 (**Figure S5G**). This binding orients BACH1 so that its lysine-rich DNA-binding domain is placed near the catalytic center of the second CUL2^FEM1B^ unit (**Figures S5H**), consistent with Lys residues in the DNA-domain being prominently ubiquitylated in cells ^63^ and potentially explaining why E3 ligase dimerization promotes BACH1 degradation.

Most importantly, our structure revealed how heme stabilizes the E3 ligase-substrate complex. In a manner not seen for other compound-induced protein interactions, heme engages three polypeptides at a time by forming interfaces with both subunits of the BACH1^CT^ dimer as well as one FEM1B molecule (**Figure 3G**). At one extended interface, heme is sandwiched between two BACH1^CT^ protomers (**Figure 3I**). The central heme iron is coordinated by the Cys646 residues of each BACH1 subunit, while the periphery of the porphyrin is engaged by the Cys621 and Cys625 residues of each BACH1. This mode of heme binding is expected to stabilize the BACH1^CT^ dimer.

In addition, the carboxy group of the heme porphyrin engages Lys16 of FEM1B and thus directly recognizes the E3 ligase (**Figure 3J**). Our structure therefore suggested that heme acts as a molecular glue with a hitherto unrecognized capacity: rather than stabilizing a binary interaction, it bridges three polypeptides to assemble a higher-order complex. We refer to such compounds as ternary molecular glues (**Supplementary Movie 1**).

To provide initial validation for this structure, we assessed the effects of disrupting each interface onto BACH1 stability. The Cys646 residue in BACH1 that coordinates the heme iron, the Cys621/Cys625 residues that clamp the porphyrin, and iron itself were all required for BACH1 turnover (**Figure 3K; Figure S2H; Figure S5I, J**); this underscored that heme-binding to BACH1 promotes degradation. Mutation of L630 and Q634 at the BACH1^CT^ dimer interface disrupted BACH1 degradation (**Figure S5G, K**); this shows that BACH1^CT^ must dimerize for turnover. L617 and L620 in the BACH helix that docks against FEM1B as well as the FEM1B residues T19, A22, or L18/L25 that bind the BACH1 helix were also required for heme-dependent degradation (**Figure 3L, M; Figure S5L**). Moreover, mutation of Y461 in the second BACH1^CT^ subunit or K60 in FEM1B stabilized BACH1 (**Figure S5M, N**); together, this confirms that the BACH1^CT^ dimer engages two sites on FEM1B to drive BACH1 degradation. BACH1 was also stabilized by mutation of FEM1B-K16, which docks against the porphyrin carboxy-group (**Figure 3N**), validating the second heme interface in the complex. Combined mutation of K16, L18, and L25 in FEM1B to disrupt docking of both heme and the heme-stabilized BACH1^CT^ dimer completely shut off BACH1 turnover (**Figure 3O**). By contrast, mutations in the canonical substrate-binding groove in FEM1B (Zn^2+^-interface: C186; R126 pocket: W93, E102, F130; aromatic pocket: V391/Q394, F501/H502) did not affect BACH1 turnover in response to heme (**Figure S5O**). We conclude that heme directs BACH1 to the same surface in FEM1B that binds the mitochondrial anchor TOM20. To accomplish this regulation, heme must interact with three polypeptides at a time and hence may act as a ternary molecular glue. Notably, the FEM1B residues that contact BACH1 are not conserved in FEM1A or FEM1C (**Figure S6A**), explaining the selectivity of CUL2^FEM1B^ in exerting this unusual proteolytic control.

### Heme acts as a ternary molecular glue

As glues typically stabilize interactions between two proteins, we wished to directly test whether heme indeed has the intriguing ability of a small molecule to act as a ternary molecular glue. We therefore established a biolayer interferometry approach to monitor the interaction between recombinant BACH1^CT^ and FEM1B. We found that BACH1^CT^ engaged FEM1B in the presence of heme with an apparent K_D_ of 0.7µM (**Figure 4A; Figure S6B-E**), while only background binding was detected in the absence of heme (**Figure 4B; Figure S6C**). This confirms that heme is required for the direct binding of BACH1 to FEM1B. Subsequent titrations showed that heme induced the association of BACH1^CT^ with FEM1B in a dose-dependent manner (**Figure 4C**), acting in part by slowing complex dissociation as previously seen with molecular glues ^64,65^. We validated heme’s ability to elicit dose-dependent binding of BACH1^CT^ to FEM1B using an orthogonal pulldown strategy (**Figure S6F**) and showed that the same heme concentrations that promoted complex formation triggered BACH1 ubiquitylation through CUL2^FEM1B^ (**Figure 4D; Figure S6G**). Heme therefore stimulates a direct interaction between FEM1B and BACH1 that results in ubiquitylation, consistent with its proposed role as a ternary molecular glue.

**Figure 4:**
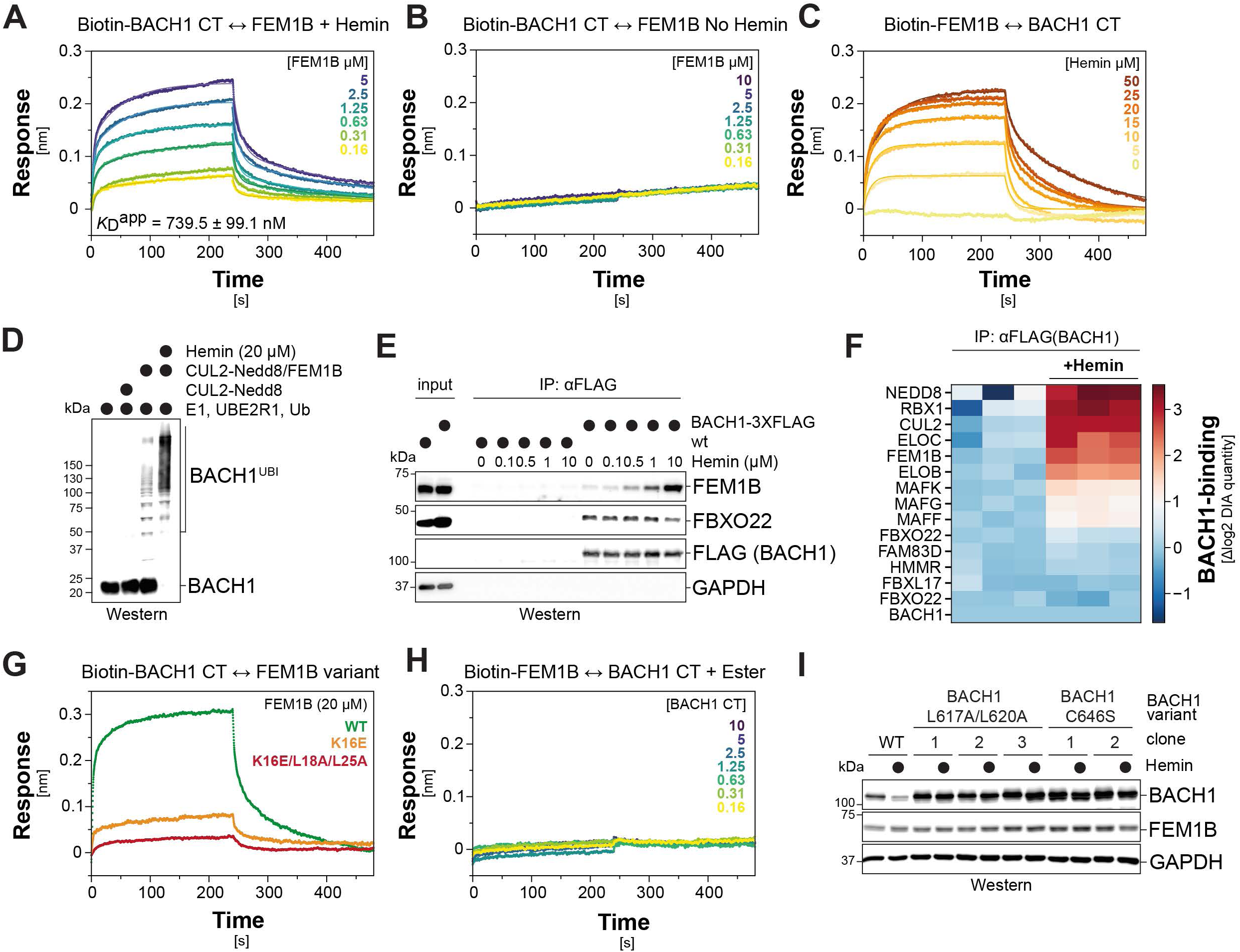
Heme acts as a ternary molecular glue. **A.** BACH1^CT^ directly binds FEM1B in the presence of hemin, as seen by biolayer interferometry. Streptavidin tips coupled to ^Biotin^BACH1^CT^ were incubated with increasing concentrations of FEM1B in the presence of 20µM hemin and association and dissociation were monitored over time. Sensorgrams were fit to a 2:1 heterogeneous ligand binding model. Fitted curves are overlaid with experimental data. The apparent K_D_ in the presence of 20 µM hemin was determined using steady-state analysis shown in Figure S6D. Similar results were obtained in n=2 independent experiments. **B.** BACH1^CT^ does not bind to FEM1B in the absence of hemin, as seen by biolayer interferometry. **C.** Heme induces binding of BACH1^CT^ to FEM1B in a dose-dependent manner. Streptavidin tips coupled to ^Biotin^FEM1B were incubated with saturating concentrations of BACH1^CT^ (20µM) and increasing concentrations of hemin. Association and dissociation were monitored over time. Similar results were obtained in n=2 independent experiments. **D.** Efficient ubiquitylation of BACH1^CT^ by CUL2^FEM1B^ requires heme. *In vitro* ubiquitylation of recombinant BACH1^CT^ by NEDD8-modified CUL2 or CUL2^FEM1B^, E1, UBE2R1, and ubiquitin was analyzed by Western blotting with an antibody that recognizes an epitope within BACH1^CT^. Similar results were obtained in n=5 independent experiments. **E.** Interaction of endogenous BACH1 with FEM1B depends on heme. HEK293T cells expressing BACH1^3XFLAG^ from the endogenous *BACH1* loci were lysed in the presence of increasing concentrations of hemin and subjected to immunoprecipitation. Co-purifying proteins were detected by Western blotting with the indicated antibodies. Similar results were obtained in n=2 independent experiments. **F.** Heme selectively induces binding of BACH1 to CUL2^FEM1B^. BACH1-3XFLAG was stably expressed in MV4-11 cells and immunoprecipitated with or without 10µM hemin. Binding partners were determined by mass spectrometry. Heatmap depicts log_2_ fold change compared to average of CTRL samples of n=3 samples. **G.** K16 in FEM1B is required for heme-dependent recognition of BACH1. Streptavidin tips coupled to Biotin-BACH1^CT^ were incubated with 20µM FEM1B variants and 20µM hemin. Association and dissociation were monitored by biolayer interferometry over time. Similar results were obtained in n=2 independent experiments. **H.** Carboxy-groups are required for heme to mediate binding of BACH1 to FEM1B. Streptavidin tips coupled to Biotin-FEM1B were incubated with increasing concentrations of BACH1^CT^ in the presence of 20µM Fe Protoporphyrin IX dimethyl ester. Similar results were obtained in n=2 independent experiments. **I.** The heme-dependent interface between FEM1B and BACH1 is required for degradation of endogenous BACH1. L617A/L620 and C646 in the endogenous *BACH1* loci of MOLM13 cells mutated using CRISPR/Cas9-mediated genome engineering. Cells were treated with 10µM hemin for 16h and lysates analyzed by Western blotting using the indicated antibodies. Similar results in n=2 independent experiments.

Immunoprecipitations showed that heme also recruited endogenous BACH1 to FEM1B in a dose-dependent manner, while FBXO22 engaged BACH1 independently of heme (**Figure 4E; Figure S6H, I**). As shown by detection of either heme or iron, heme was a central component of these BACH1-FEM1B assemblies (**Figure S6J, K**). Mass spectrometry analyses revealed that heme almost exclusively stabilized the association of BACH1 with subunits of the CUL2^FEM1B^ E3 ligase, while other interactors, including MAF transcription factors, FBXO22, or FBXL17, were only modestly affected or did not change at all (**Figure 4F**). These findings underscored that heme specifically recruits BACH1 to CUL2^FEM1B^, which is also in line with an activity as a molecular glue.

Having reconstituted the heme-dependent recognition of BACH1 by FEM1B, we assessed the effects of mutations or chemical alterations onto complex formation. Mutation of FEM1B-K16, which contacts a carboxy-group of the heme porphyrin, strongly impaired binding to BACH1^CT^ in biolayer interferometry (**Figure 4G**). Additional mutation of L18 and L25 in FEM1B to disrupt recognition of the heme-stabilized BACH1^CT^ dimer, abrogated any residual interaction (**Figure 4G; Figure S6L**). Moreover, replacing the heme carboxy-group with an uncharged ester that fails to engage FEM1B-K16 abolished its ability to stabilize the BACH1-FEM1B complex (**Figure 4H; Figure S6M**). Mutation of the heme-coordinating C646 in BACH1 also blocked its heme-depen-dent binding to FEM1B (**Figure S6N**). We conclude that heme must contact both FEM1B and BACH1 to promote their interaction, providing strong evidence for a role as ternary molecular glue that bridges three polypeptides for the assembly of a higher-order E3 ligase-substrate complex.

To assess the ternary glue mechanism in cells, we generated variants impaired in heme recognition (FEM1B^K16E^; BACH1^C646S^; BACH1^C621S/C625S^) or heme-dependent complex formation (FEM1B^L18A/L25A^; FEM1B^T19E^; FEM1B^A22E^; BACH1^L617A/L620A^; BACH1^Y641A^). All mutations that blocked ternary glue function *in vitro* abolished heme-dependent interactions in cells (**Figure S7A, B**). Proteomic analyses showed that impeding heme binding disrupted the interaction of BACH1 with endogenous CUL2^FEM1B^, but barely any other protein (**Figure S7C**). To assess consequences for endogenous interactions, we introduced the C646S or L617A/L620A mutations into the *BACH1* loci of two AML cell lines (**Figure S7D**). These mutations increased the abundance of endogenous BACH1 under basal conditions and resulted in a protein that was no longer turned over upon hemin treatment (**Figure 4I; Figure S7E**). We conclude that heme acts as a ternary molecular glue that selectively recruits BACH1 to CUL2^FEM1B^ and thereby induces the proteasomal degradation of the transcriptional repressor.

### Ternary glue signaling enables localized stress sensing

As ternary glues had not been observed before, we next wished to determine the biological out-puts resulting from such signaling. Given that BACH1 is a transcriptional repressor, we hypothe-sized that its degradation might elicit the expression of genes that allow cells to respond to altered heme levels. To prevent secondary effects of prolonged BACH1 modulation, we acutely depleted either BACH1 to model degradation or FEM1B to generate cells unable to rapidly eliminate BACH1 in response to heme. Using RNA sequencing, we found that only ∼80 genes were induced upon acute loss of BACH1, but repressed if cells were depleted of FEM1B (**Figure 5A, B**). The single most strongly co-regulated gene was *HMOX1*, which encodes an oxygenase that degrades labile heme in the cytoplasm (**Figure 5A, B**). Quantitative RT-PCR analyses confirmed that loss of FEM1B reduced *HMOX1* expression, while BACH1 depletion strongly upregulated this gene (**Figure 5C; Figure S8A, B**). As co-depletion of FEM1B and BACH1 restored HMOX1 mRNA to control levels, CUL2^FEM1B^ and BACH1 oppose each other in regulating HMOX1 expression (**Figure 5C**). This regulation is direct, as chromatin immunoprecipitations showed that CUL2^FEM1B^ restricted the recruitment of BACH1 to the *HMOX1* enhancers (**Figure 5D; Figure S8C**). These results indicated that acute degradation of BACH1 primarily elicits the production of HMOX1, an enzyme that protects cells from excessive heme ^66^.

**Figure 5:**
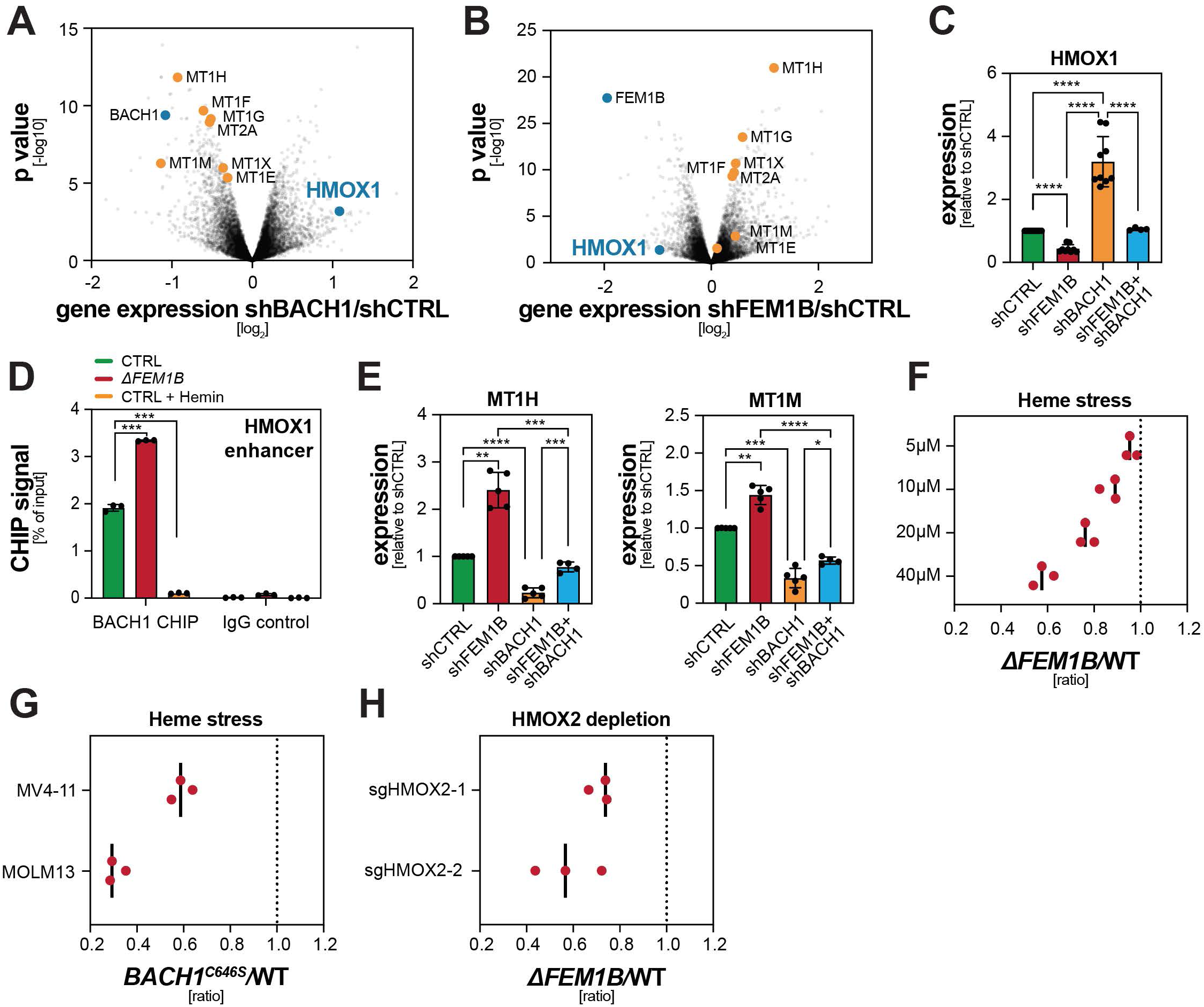
Ternary glue signaling elicits a gene expression response to alleviate heme toxicity. **A.** RNA sequencing analysis reveals genes under control of BACH1. MV4-11 cells were depleted of BACH1 using specific shRNAs. RNA was purified and analyzed by global RNA sequencing. Differentially expressed genes are depicted as volcano plots. n=3 independent experiments. **B.** RNA sequencing analysis reveals genes under control of FEM1B. MV4-11 cells were depleted of FEM1B using specific shRNAs. RNA was purified and analyzed by global RNA sequencing. Differentially expressed genes are depicted as volcano plots. n=2 independent experiments. **C.** BACH1 and FEM1B co-regulate HMOX1 expression. BACH1 and/or FEM1B were depleted in MV4-11 cells using specific shRNAs, and HMOX1 expression was analyzed by qPCR. Data of n=4-9 independent experiments represented as mean ± SD. **D.** FEM1B restricts BACH1 binding to the HMOX1 enhancer. Chromatin immunoprecipitations of endogenous BACH1 from WT or *ΔFEM1B* MV4-11 cells were analyzed by qPCR, using primers directed against validated BACH1 binding sites in the EN2 enhancer of HMOX ^87^. Cells treated with hemin to induce degradation of BACH1 were used as control. Data expressed as percentage of input. Similar results observed in n=3 independent experiments. **E.** BACH1 and FEM1B both impact metallothionein transcription. BACH1 and/or FEM1B were depleted in MV4-11 cells using specific shRNAs and expression of MT1H or MT1M were monitored by qPCR. Data of n=4-5 independent experiments represented as mean ± SD. **F.** *ΔFEM1B* AML cells are sensitive to heme toxicity. mCherry-expressing MV4-11 *ΔFEM1B* cells were mixed with GFP-expressing WT cells and mCherry/GFP ratios were determined by flow cytometry after 4d of co-culture in the presence of increasing hemin concentrations. n=3 independent experiments. **G.** CUL2^FEM1B^-resistant *BACH1^C646S^*::cells are sensitive to heme. mCherry-expressing BACH1^C646S^ MV4-11 or MOLM13 cells were co-cultured with corresponding WT cells in the presence or absence of 40µM hemin and analyzed as above. n=3 independent experiments. **H.** *ΔFEM1B* cells are sensitive to increased endogenous heme upon depletion of the constitutively expressed heme-degrading enzyme HMOX2. WT (GFP) and *ΔFEM1B* (mCherry) MV4-11 cells were mixed, depleted of HMOX2 and mCherry/GFP ratios were determined after 12d of competition. n=3 independent experiments. Statistical significance in (C), (D) and (E) was determined using one-sample t-tests and two-tailed Student’s t-tests (* p<0.1, ** p<0.01, *** p<0.001 and **** p<0.0001).

While expression of HMOX1 can be induced by NRF2 ^67^, other targets of this transcription factor, including regulators of the ETC or ferroptosis ^53,68^, were not co-regulated by acute BACH1 or FEM1B depletion. Dysregulated ternary glue signaling is therefore unlikely to simply induce oxidative stress. Instead, the RNAseq and qRT-PCR analyses revealed that CUL2^FEM1B^ inhibition and subsequent BACH1 stabilization caused expression of cysteine-rich metallothionein proteins that scavenge prooxidant metal ions and ROS molecules ^69^ (**Figures 5B, 5E; Figure S8D**). Loss of BACH1, which induces HMOX1 to degrade heme, had the opposite effect and resulted in low metallothionine mRNA levels (**Figure 5A, 5E; Figure S8D**). These observations suggested that acute BACH1 degradation through a ternary glue allows cells to respond to a specific stress that is caused by excess labile heme.

Notably, heme elicits stress in a localized manner: while cytoplasmic free heme causes stress, mitochondrial heme is destined for incorporation into ETC complexes and thereby provides cells with an important function. As the ternary glue tethers BACH1 to the same surface on FEM1B that is also engaged by TOM20, only cytoplasmic CUL2^FEM1B^ should degrade BACH1, and cells should therefore only convert cytoplasmic, but not mitochondrial, heme accumulation into a stress response. Several lines of evidence supported the notion that TOM20-associated, mitochondrial CUL2^FEM1B^ is unable to drive BACH1 turnover: recombinant TOM20 competed BACH1 off FEM1B in a dose-dependent manner (**Figure S8E**); TOM20 strongly impaired BACH1 ubiquitylation by CUL2^FEM1B^ (**Figure S8F**); and TOM20 overexpression stabilized BACH1 in cells (**Figure S8G**). Moreover, tethering CUL2^FEM1B^ to mitochondria blocked BACH1 degradation, while it did not affect the turnover of a canonical substrate, FNIP1 (**Figure S8H**). Thus, BACH1 can only be degraded via CUL2^FEM1B^ complexes that do not associate with mitochondria. Ternary glue signaling there-fore ensures that CUL2^FEM1B^ senses heme in the cytoplasm - where it causes stress ^70^ - while ignoring mitochondrial heme that may transiently accumulate prior to its incorporation into the ETC. In essence, ternary glues establish a mechanism of localized signaling through a small molecule.

To test if localized stress signaling preserves cell fitness, we exposed mixtures of WT and *ΔFEM1B* cells to increasing concentrations of hemin. We found that *ΔFEM1B* AML cells were highly sensitive to hemin (**Figure 5F**). The same phenotype was observed in cells that expressed endogenous BACH1^C646S^, showing that it is ternary glue signaling that protects cells from stress induced by excess free heme (**Figure 5G**). *ΔFEM1B* AML cells also showed reduced fitness when depleted of HMOX2 (**Figure 5H**), a constitutively expressed oxygenase that selectively degrades cytosolic heme ^71^. We conclude that ternary glue signaling constitutes a stress response that protects cells from accumulation of toxic cytoplasmic heme. By ensuring that CUL2^FEM1B^ senses heme away from mitochondria, cells avoid triggering this response during times of ETC-production. The ternary glue mechanism therefore integrates localization of a small molecule with its effector function to orchestrate signaling beyond simple stabilization of protein interactions.

### Loss of FEM1B sensitizes AML cells to the BCL2-inhibitor Venetoclax

The importance of ternary glue signaling for AML fitness showed that these cancer cells must limit accumulation of labile heme in the cytoplasm. Supporting this notion, we noted in transcriptomic analyses of bone marrow aspirates of 675 cancer patients that AML cells downregulate the heme biosynthesis enzymes ALAS1, ALAD, HMBS, UROD, UROS, CPOX, and FECH compared to un-transformed controls (**Figure 6A**). Reduced expression of these enzymes was observed across all genetic AML subtypes (**Figure 6B; Figure S9A**), consistent with metabolic rewiring occurring independently of genotype ^32^. These observations were in line with data from other patient cohorts^42^ and implied that disrupting ternary glue signaling by inhibiting CUL2^FEM1B^ could offer opportunities for AML treatment.

**Figure 6:**
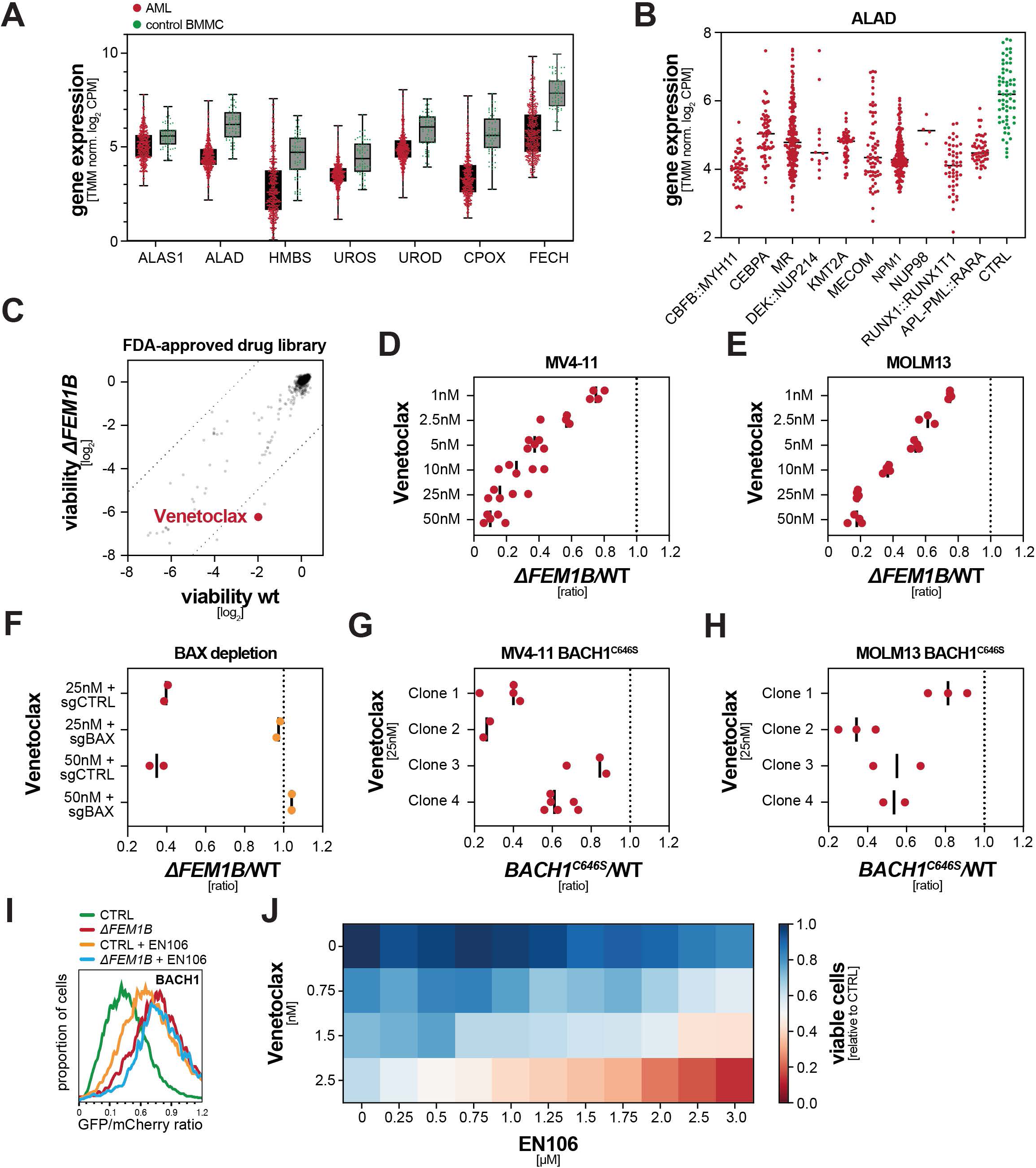
Disrupting ternary glue signaling sensitizes AML cells to the BCL2-inhibitor Venetoclax. **A.** Expression of heme biosynthesis enzymes is downregulated in AML patients, as shown by gene expression analysis in mononuclear cells derived from bone marrow aspirates of 675 patients compared to bone marrow mononuclear cells (BMMC) of 64 healthy controls. TMM-normalized log_2_CPM values expressed as individual datapoints and boxes (median and 25^th^/75^th^ percentile) and whiskers (min to max). **B.** Expression of the heme biosynthesis enzyme ALAD is downregulated in all AML subtypes. Gene expression analysis of patients and controls described above separated by genetically defined subtypes according to the WHO 2022 classification. Individual datapoints shown together with mean. **C.** Synthetic lethality drug screen reveals that *ΔFEM1B* AML cells are hypersensitive to Venetoclax. WT and *ΔFEM1B* MV4-11 cells were treated with 1200 FDA-approved compounds at 1µM for 48h, after which cell viability was determined using Cell TiterGlo. **D-E.** Cell competition confirms strong synergy between *FEM1B* deletion and Venetoclax treatment. mCherry-expressing *ΔFEM1B* and GFP-expressing WT MV4-11 (D.) or MOLM13 (E.) cells were mixed and incubated with the indicated concentrations of Venetoclax for 4d. mCherry/GFP ratios were determined by flow cytometry. n=3-5 independent experiments. **F.** Depletion of pro-apoptotic BAX rescues synthetic lethality of combined BCL2- and FEM1B-inhibition. mCherry-expressing *ΔFEM1B* and GFP-expressing WT MV4-11 cells were infected with sgRNAs targeting BAX, mixed, and treated with 25nM or 50nM Venetoclax for 4d. mCherry/GFP ratios were determined by flow cytometry. n=2 independent experiments. **G-H.** Heme-resistant BACH1^C646S^ cells are hypersensitive to Venetoclax treatment. mCherry-expressing endogenously edited BACH1^C646S^ MV4-11 (G.) and MOLM13 (H.) cells were mixed in equal numbers with GFP-expressing WT cells and incubated with 25nM of Venetoclax for 4d. mCherry/GFP ratios were determined by flow cytometry. n=2-6 independent experiments. **I.** BACH1 is stabilized by inhibition of CUL2^FEM1B^ with EN106. BACH1 stability reporters were transfected into wt and *ΔFEM1B* HEK293T cells, which were then treated with 20µM EN106 for 16h. BACH1 stability was monitored by flow cytometry. Similar results in n=2 independent experiments. **J.** Venetoclax and EN106 show synthetic lethality in AML cancer cells. MV4-11 cells were treated with increasing concentrations of Venetoclax and EN106 and viability was determined by Cell TiterGlo after 48h. Data points represent the mean of 3 technical replicates. Similar results observed in n=3 independent experiments.

To convert this metabolic vulnerability into an effective therapeutic strategy, we exploited a chemical synthetic lethality screen design ^72^ to search for FDA-approved agents that are selectively toxic in combination with CUL2^FEM1B^ inhibition. We found that loss of *FEM1B* strongly synergized with Venetoclax (**Figure 6C**), a BCL2 inhibitor that has been approved in AML in combination with hypomethylating agents or cytarabine ^73^. Cell competition assays confirmed the synergy between *FEM1B* loss and Venetoclax treatment in several AML lines (**Figure 6D, E; Figure S9B**). Similar to *ΔFEM1B* cells, acute FEM1B depletion caused a striking synthetic lethality with Venetoclax treatment (**Figure S9C, D**), which was also observed with other BCL family inhibitors (**Figure S9E-H**). Combined FEM1B- and BCL2-inhibition specifically induced apoptosis (**Figure S9I**), which could be overcome by co-depletion of pro-apoptotic BAX (**Figure 6F**) or treatment with the caspase inhibitor Z-VAD-FMK (**Figure S9J**). Most importantly, cells that expressed BACH1^C646S^ from the endogenous *BACH1* loci were also highly sensitive to Venetoclax (**Figure 6G, H**), showing that it is defective ternary glue signaling that sensitizes AML cells to Venetoclax treatment.

These observations encouraged us to directly test the effects of combined small molecule inhibition of CUL2^FEM1B^ and BCL2 in AML cells. Having developed an early covalent CUL2^FEM1B^ inhibitor, EN106 ^74^, we first showed that this compound stabilized BACH1 (**Figure 6I**). Importantly, exposing AML cells to increasing concentrations of EN106 revealed a marked synthetic lethality with Venetoclax (**Figure 6J**). While EN106 is a tool compound, these findings point to combination treatment with CUL2^FEM1B^- and BCL2-inhibitors for aggressive AML subtypes, showing that the discovery of ternary glue signaling rapidly informs therapeutic strategies in a cancer of high unmet need.

## Discussion

Both therapeutic molecules and endogenous metabolites can act as molecular glues that stabilize protein interactions to rewire crucial signaling networks. While known glues have all been reported to constitutively promote binary interactions, cells use a distinct mechanism to detect and degrade excess heme: heme bridges three polypeptides to tether the transcriptional repressor BACH1 to the mitochondrial-binding motif of the E3 ligase CUL2^FEM1B^, thereby ensuring that only cytoplasmic heme elicits BACH1 degradation to induce expression of HMOX1. Heme is a therefore a ternary molecular glue, a hitherto unrecognized class of small molecule that can integrate multiple inputs - here, heme stress and compound localization - for more precise biological outputs (**Figure 7**). This work therefore provides a blueprint for developing small molecules that exert their functions in a localized manner.

**Figure 7:**
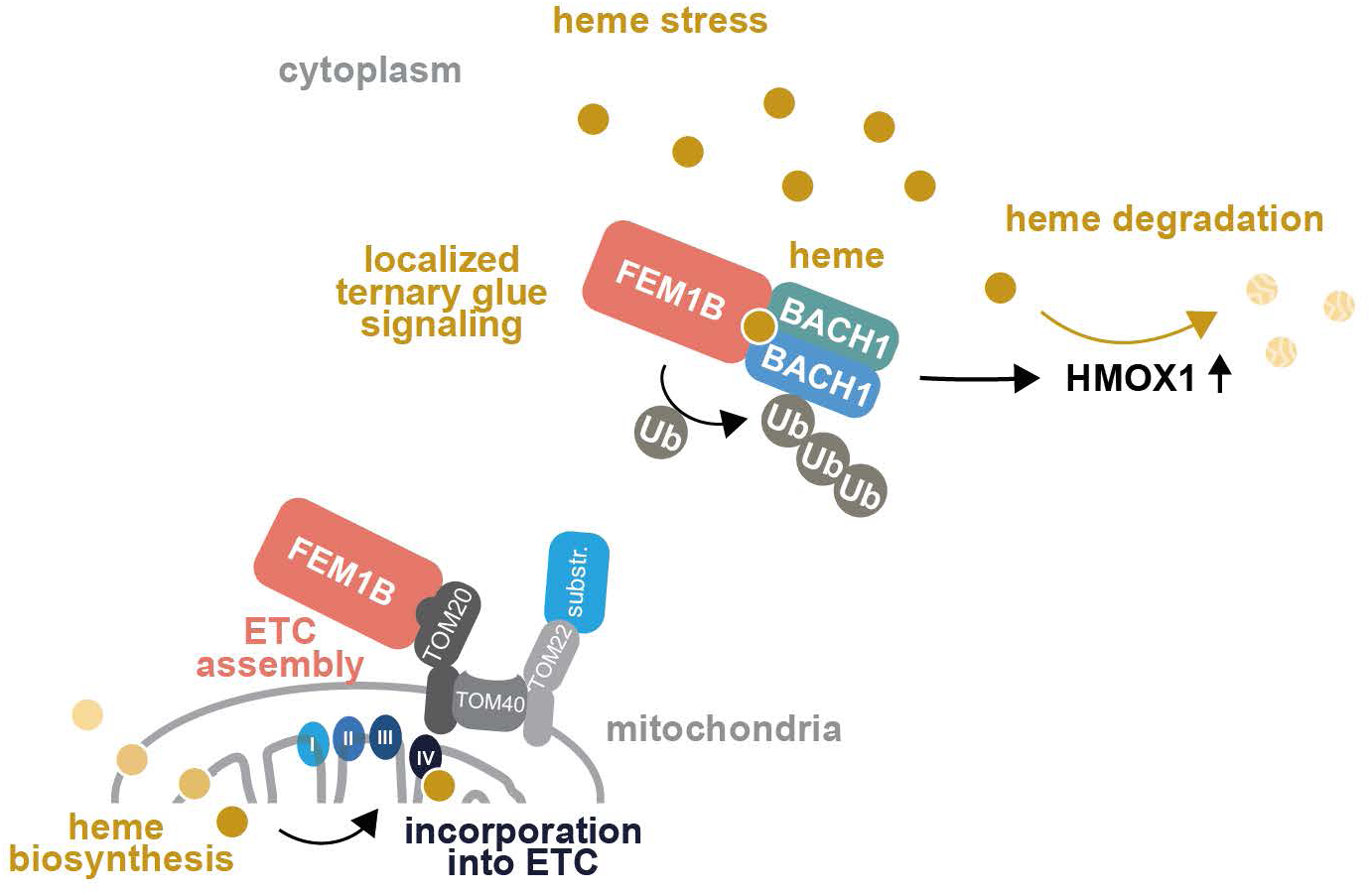
The ternary glue mechanism establishes localized signaling by a small molecule. CUL2^FEM1B^ senses excess heme in the cytoplasm, where it causes stress, while ignoring mito-chondrial heme destined for incorporation in the ETC. To accomplish this regulation, heme acts as a ternary glue that tethers a BACH1 dimer to the same site in FEM1B that is also used to recruit the E3 ligase to mitochondria. In this manner, heme can only induce BACH1 degradation through cytosolic CUL2^FEM1B^ complexes that are not associated with mitochondria, thus selectively eliciting expression of the heme-degrading enzyme HMOX1 when prooxidant heme accumulates in the cytoplasm.

Molecular glues typically act in a cooperative fashion by strengthening weak preexisting interactions between proteins with complementary surfaces ^3^. Rapamycin, FK506, IMiDs, indisu-lam, UM171, plant hormones, or purines accordingly bridge two proteins. Heme functions in a strikingly different manner by engaging three polypeptides at a time. It induces dimerization of the carboxy-terminal BACH1 domain by sandwiching in-between the Cys/Pro motifs of each subunit, and it docks this BACH1^CT^ dimer to a surface of FEM1B that is used to anchor the E3 ligase to mitochondrial membranes. Other features of molecular glues are preserved: the surfaces bet-ween partners are complementary to allow the BACH1 dimer to engage two proximal sites on FEM1B. Moreover, all interfaces between heme, FEM1B, and BACH1 are required for BACH1 turnover, suggesting that complex formation may occur in a cooperative manner. While our discovery of a ternary glue reveals that induced proximity molecules can act beyond dimerization, other previously defined rules of glue function still apply.

The mechanism of heme ternary glue signaling ensures that cells selectively sense heme away from mitochondria, where the final step of heme synthesis is carried out and where heme is further modified for incorporation into cIV. The heme ternary glue is a therefore a small molecule that acts in a localized manner. It is possible that cytoplasmic CUL2^FEM1B^ is simply more efficient in targeting a transcriptional repressor that does not possess mitochondrial targeting sequences and hence does not efficiently reach this organelle. We consider it more likely, however, that localized BACH1 degradation isolates stress signaling from sites of heme function in the ETC, a metabolic pathway that is particularly important in AML cells ^30,75–80^. Impaired ETC assembly can lead to transient accumulation of heme in the mitochondrial membrane ^81,82^, and if these con-ditions persist, these heme molecules diffuse into the cytoplasm to exert stress. Strikingly, ETC inactivation turns on CUL2^FEM1B^ ^35–37^, suggesting that the same E3 ligase alleviates mitochondrial stress by inducing cIV assembly and counteracts cytoplasmic stress arising from impaired heme incorporation. The ternary glue mechanism therefore allows cells to address multiple facets of ETC stress through a single effector that processes key substrates at distinct sites. Molecular glues can therefore regulate signaling much more precisely than previously appreciated.

Our findings have important implications for the treatment of AML, a cancer whose genetic heterogeneity has complicated patient stratification for effective therapy. Recent work showed that aggressive AML subtypes depend on high mitochondrial activity and rapid ETC assembly inde-pendently of genotype ^32^. Proliferation of such AML cells was strongly impaired by *FEM1B* deletion, a condition that at the same time impedes cIV assembly and prevents clearance of pro-oxidant heme. This metabolic vulnerability can be exacerbated by co-treatment with Venetoclax, a BCL2 inhibitor that is already in use for AML therapy. Venetoclax-treated cells that lack *FEM1B;* express a BACH1 variant resistant to heme-induced degradation; or face the CUL2^FEM1B^ inhibitor EN106 undergo apoptosis much more readily than cells that can degrade BACH1. As AML cells depend on constant heme synthesis and ETC assembly, disrupting ternary glue signaling will likely affect cancer cells more drastically than their untransformed counterparts, thus hopefully overcoming the toxicity associated with targeting the core ETC machinery.

Inspired by the mechanism of purine and heme sensing ^21^, we expect that discovering endogenous molecular glues will continue to inform therapeutic approaches for diseases with high unmet need. Our identification of heme as a ternary glue shows that induced proximity molecules can act beyond simple complex formation and integrate additional information about the cellular state: by anchoring BACH1 to a regulatory site in FEM1B, cells use a small molecule to trigger a localized stress response. As prospective glue discovery has become a possibility ^15,16^, designing screens to direct proteins to regulatory sites in effectors that control localization, activation ^37,83,84^, or stability ^85^, could tailor the site, strength, or duration of the drug effect to needs defined by a disease. Thus, uncovering physiological mechanisms of glue signaling will help expand the reach of induced proximity as a therapeutic modality.

## Acknowledgements

We thank all of the members of our laboratories, especially D. Haakonsen, for enthusiastic support and many suggestions for this work; J. Zuber (IMP Vienna) for sharing the human whole genome sgRNA library and R. Kalis and F. Andersch for help with data processing; QB3 Genomics, UC Berkeley, Berkeley, CA, for Illumina sequencing; K. Sharma, D. Toso and the members of the Cal-Cryo cryo-EM facility at QB3-Berkeley for assistance on cryo-EM data collection; the Cancer Research Laboratory Flow Cytometry Facility; the Inductively Coupled Plasma Spectroscopy Facility; the UCB Cell Culture Facility which is supported by The University of California Berkeley (RRID SCR_017924); and M. Ghassemian and the members of the UCSD Biomolecular and Proteomics Mass Spectrometry facility for assistance in running proteomics MS samples. M.H. was funded by the Deutsche Forschungsgemeinschaft (DFG, German Research Foundation; HE 9330/1-1), R.S. by the American Heart Association (24PRE1195753). This work was supported by the Howard Hughes Medical Institute (M.R), an NIGMS RO1 to MR (GM151335-01) and the Molecular Therapeutics Initiative (J.S. and E.W.). M.R. is an Investigator of the Howard Hughes Medical Institute.

## Contributions

M.H. performed experiments and analyses including CRISPR screening, proteomics, cell culture experiments with help from C.H., H.X., T.B., R.S. and S.C.. C.H. and M.H. performed in vitro studies. Z.Y. collected and processed cryo-EM data. W.W. and T.H. provided and analyzed patient datasets. E.W. and J.S. designed, performed and analyzed the small molecule synthetic lethality screen. M.R. helped to plan and interpret experiments. M.H. and M.R. wrote the manuscript with input from all authors.

## STAR Methods

### EXPERIMENTAL MODEL AND STUDY PARTICIPANT DETAILS

#### Cell lines

Human embryonic kidney (HEK) 293T cells were maintained in DMEM + GlutaMAX (Gibco, 10566-016), MV4-11 cells were maintained in IMDM + GlutaMAX(Gibco, 31980-030), MOLM13 and THP1 cells were maintained in RPMI 1640 (Gibco, 11875-093), all supplemented with 10% fetal bovine serum (VWR, 89510-186) and penicillin/streptomycin (Gibco, 15140-122). Expi293F cells were maintained in Expi293™ Expression Medium (Gibco, A1435101) in shaker flasks. All cells were purchased from the Berkeley Cell Culture Facility (authenticated by short tandem repeat analysis). All cell lines were routinely tested for mycoplasma contamination using the Mycoplasma PCR Detection Kit (abmGood, G238). All cell lines tested negative for mycoplasma. Plasmid transfections for were performed using polyethylenimine (PEI) at a 1:6-1:8 ratio of DNA (in mg) to PEI (in ml at a 1 mg/ml stock concentration). Lentiviruses were produced in HEK 293T cells by co-transfection of lentiviral and packaging plasmids using TransIT®-Lenti Transfection Reagent (Mirus, MIR 6603) according to the manufacturer’s protocol. Viruses were harvested 72 h post transfection, aliquoted, and stored at 80°C for later use. For lentiviral transduction, 10^5^ of HEK293T cells were seeded into 24-well plates or 5x10^5^ AML cells were seeded into 12-well plates. Cells were subjected to centrifugation for 30 min at 700g after addition of lentiviral particles and 8 μg/ml polybrene (Sigma-Aldrich, TR-1003). Transduced cells were drug-selected 24 h after infection with the following drug concentrations: puromycin (HEK293T, THP1: 1 μg/ml; MV4-11, MOLM13: 0.5µg/ml Sigma-Aldrich, P8833); blasticidin (all cell lines: 10 μg/ml; Thermo Fisher, A1113903). Recombinant proteins were produced in were expressed in LOBSTR-BL21(DE3)-RIL cells (Vector Laboratories, NC1789768) grown in LB broth media.

#### Human studies

The study was conducted in accordance with the Declaration of Helsinki and the Ethics Committee of the Bavarian Medical Association for the use of archived RNA samples and associated clinical information. The clinical data were retrieved, and the samples were collected and analyzed with the endorsement of the Ethics Committee of the Bavarian Medical Association. All patients had given a written informed consent to the use of genetic and clinical data according to the Declaration of Helsinki.

Library preparation was done as previously described ^90^. In brief, total RNA were extracted from lysed cell pellet of diagnostic bone marrow. Two hundred fifty ng of high-quality RNA were used as input for the TruSeq Total Stranded RNA kit (Illumina, San Diego, CA, USA). 101bp paired-end reads were produced on a NovaSeq 6000 system with a median yield of 54 million cluster per sample. FASTQ generation was performed applying Illumina’s bcl2fastq software (v2.20). Using BaseSpace’s RNA-seq Alignment app (v2.0.1) with default parameters, reads were mapped with STAR aligner (v2.5.0a, Illumina) to the human reference genome hg19 (RefSeq annotation). Estimated read counts per gene were obtained from Cufflinks 2 (version 2.2.1). Non expressed genes were filtered out (<2 counts). Raw counts were normalized by applying the Trimmed mean of M-values method from the edgeR package ^91^, producing log2 CPM values. Expression data can be found in Supplementary Table 5.

## METHOD DETAILS

### Antibodies

The following antibodies were used in this study: Rabbit monoclonal DYKDDDDK Tag Antibody (Cell Signaling Technology, 2368), mouse monoclonal anti-Flag clone M2 (Sigma-Aldrich, F1804), rabbit monoclonal anti-GAPDH (14C10) (Cell Signaling Technology, 2118S), rabbit polyclonal anti-CUL2 (Bethyl, A302-476A), rabbit monoclonal anti-FNIP1 (Abcam, ab134969), rabbit polyclonal anti-FEM1B (Proteintech, 11030-1-AP), rabbit polyclonal anti-BACH1 (Proteintech, 14018-1-AP), rabbit monoclonal anti-BACH1 (E4E7B, for CHIP, Cell Signaling Technology #33059), mouse monoclonal anti-BACH1 (F-9, Santa Cruz Biotechnology, sc-271211), normal rabbit IgG (Cell Signaling Technology #2729), rabbit polyclonal anti-FBXO22 (Proteintech, 13606-1-AP), mouse monoclonal anti-Alpha Tubulin (Sigma-Aldrich, CP06), goat anti-rabbit IgG (H+L) HRP (Vector Laboratories, PI-1000-1), sheep anti-mouse IgG (H+L) HRP (Sigma, A5906), goat anti-mouse IgG light-chain-specific HRP conjugated (Jackson Immunoresearch, 115-035-174) and APC anti-rat CD90/mouse CD90.1 (Thy-1.1) (Biolegend, 202526).

### Cloning

All genes were cloned from cDNA prepared from HEK293T or AML cells or ordered as gene blocks from IDT. The list of all constructs used in this study are provided in the Key resources table. Most cloning was performed using Gibson assembly using HIFI DNA Assembly master mix (NEB, E2621L).

### CRISPR-Cas9 genome editing

All CRISPR-Cas9 edited cell lines for this study were generated using Cas9-RNP complexes, the Lonza 4D-Nucleofector X Unit (program CM-130 for HEK293T cells or EN138 for AML cell lines) and the SF Cell Line 4D-Nucleofector ® X Kit S (Lonza, V4XC-2032). Alt-R™ CRISPR-Cas9 sgRNAs were ordered from IDT. Recombinant Cas9 protein was purified in house by the UCB QB3 MacroLab. ΔFEM1B cells were generated by excising the first exon of the FEM1B gene. Point mutations in or addition of a C-terminal 3X-FLAG tag to the endogenous BACH1 locus were achieved by homology directed repair using Alt-R™ HDR single stranded donor oligos (ssODN) by IDT. For each reaction, 2.5 µl of recombinant Cas9 (40 µM) was mixed with 1.3 µl of sgRNA (100 µM) with 10 µl SF nucleofector solution and incubated for 10 min at room temperature, after which 0.75 µl of ssODN (100 µM) was added in the case of HDR editing. 4x10^5^ cells were resuspended in 5µl SF nucleofector solution and mixed with the RNP complex. Immediately after nucleofection, cells were transferred to 12-well plates. To enhance HDR efficiency Alt-R™ HDR Enhancer V2 (IDT, 10007910) was added to AML cells at 300nM for max. 16 h, after which the media was exchanged. 72 h post nucleofection bulk editing efficiency was determined by genomic DNA extraction using QuickExtract DNA Extraction Solution (Biosearch Technologies, QE09050) and PCR. Editing efficiency was assessed by gel electrophoresis or Sanger sequencing followed by ICE (Inference of CRISPR Edits) analysis (EditCo). Cells were then subjected to single clonal selection in 96-well plates. After 10-14 days individual colonies originating from a single clone were picked and transferred to a 24-well plate. Homozygous clones were confirmed by PCR genotyping, DNA sequencing, western blot and qPCR analysis when applicable. Presumably due to toxicity of BACH1 accumulation we were only able to obtain heterozygous BACH1^C646S^ clones in the MV4-11 cell line. Bulk genetic depletions were carried out using the lentiCRISPRv2 puro (Addgene, 52961) vector. sgRNA and donor sequences are listed in Supplementary Tables 2-3.

### Whole genome CRISPR dropout screen

We followed a modified CRISPR–Cas9 screening protocol as previously described ^92,93^. Stable Cas9-expressing cells were generated by lentiviral delivery of the lentiCas9-Blast plasmid (Addgene, #52962), followed by selection with blasticidin and single cell selection in 96-well plates. Cas9 cutting efficiency was determined by lentiviral transduction of the pMCB320 plasmid (Addgene, #89359), which encodes for the fluorescent protein mCherry and an sgRNA targeting mCherry. After selection with puromycin, the fraction of mCherry+ cells was determined using flow cytometry. Clones with >90% depletion of mCherry+ cells (compared to cells infection with a non-targeting control sgRNA) and a comparable growth pattern to the parental cell line were used in the CRISPR screen. Pooled sgRNA library viruses were obtained by transfection of the human VBC 2-guide per gene library (38954 optimized sgRNAs targeting 19477 human genes constructed in the pETN lentiviral vector) into HEK293T cells together with lentiviral packaging plasmids using Mirus TransIT-293 Transfection reagent. In order to maintain a 1000X coverage throughout the screen, 2x10^8^ MV4-11/THP1 WT and ΔFEM1B cells were spinfected with the pooled sgRNA virus at a multiplicity of infection (MOI) of 0.2 with 8 μg/ml polybrene in 6-well plates. Media was exchanged and cells were pooled in large flasks after 24h and MOI was determined after 48h, after which selection with G418 (Invivogen, ant-gn-5) 800µg/ml (MV4-11) or 1000µg/ml (THP1) was performed for 6-8 days. MOI and selection efficiency were determined by flow cytometry of cells stained with APC anti-rat CD90/mouse CD90.1 (Thy-1.1) antibody (Biolegend, 202526), which recognizes a murine cell surface antigen encoded on the sgRNA library plasmids. When cells reached near 100% positivity for Thy-1.1, the screen was started by harvesting 5x10^7^ cells as d0 samples and cells were maintained by passaging a minimum of 5x10^7^ cells per condition every 72h. Additional samples were harvested on day 12 and day 21 of the screen (MV4-11) and day 21 and day 30 (THP1) to account for slower growth of THP1 cells. Genomic DNA extraction was performed using a QIAamp DNA Blood Maxi Kit (QIAGEN, 51194) according to the manufacturer’s instructions. sgRNA-encoding regions were amplified and staggered barcodes were added to the amplicon with AmpliTaq Gold™ DNA Polymerase (Thermo Fisher, 4311820) with the following primers (fw 5’ AGTATTAGGCGTCAAGGTCC 3’, rv 5’ CTCTTTCCCTACACGACGCTCTTCCGATCT-1-4bp stagger-4N barcode-TTCCAGCATAGCTCTTAAAC 3’). 250µg of genomic DNA were used as template for the first PCR step to maintain >1000X coverage. Pooled samples were cleaned up using ampure bead clean-up with 0.8× and 1.2× cut-off values (Beckman Coulter, A63881). Illumina adapters were added during a second PCR reaction with the following primers (fw 5’ CAAGCAGAAGACGGCATACGAGATAGTATTAGGCGTCAAGGTCC 3’, rv 5’ AATGATACGGCGACCACCGAGATCTACACTCTTTCCCTACACGACGCT 3’), followed by another bead cleanup. Samples were quantified by Qubit (Thermo Fisher, Q33230), underwent further QC and quantification with fragment analysis and qPCR and were sequenced at the UC Berkeley Vincent J. Coates Genomics Sequencing laboratory on a NovaSeqX. NGS data was demultiplexed using crispr-process-nf (https://github.com/ZuberLab/crispr-process-nf). Subsequently, CasTLE analysis ^86^ was run to identify top candidate genes that were synthetic lethal or protective in ΔFEM1B cells compared with WT cells. Combo CasTLE analysis was used to enhance statistical power and reliability of hits (two distinct ΔFEM1B clones in the MV4-11 screen and the THP1 screen, respectively). Mitochondrial proteins were defined as such if annotated in MitoCarta3.0 ^94^. Gene list enrichment analysis analysis of <5% FDR scoring depleted genes in the MV4-1 screen (n=504 genes) was performed using Enrichr ^88^.

### Protein stability reporter assay

The pCS2+-substrate-GFP-IRES-mCherry reporter constructs were generated as previously described ^39^. HEK293T cells were seeded in 6-well plates at a density of 300,000 cells. The next day, 50 ng of reporter plasmid and empty vector up to 500 ng total were transfected using PEI and collected for flow cytometry after 48 h. For stable reporter expression and sgRNA depletion experiments the reporter cassette was subcloned into pLVX-tetone-blasticidin, which was lentivirally transduced into cells, followed by selection with blasticidin. Bulk genetic depletions were carried out using lentiCRISPRv2 (Addgene, 52961). Following selection with puromycin, cells were plated in a 6-well plate at 200,000 cells/well. Reporter expression was started the next day by addition of Doxycyline 2 µg/ml and cells were analyzed after 48 h of induction using either a BD Bioscience LSR Fortessa, and the GFP/mCherry ratio was analysed using FlowJo.

### Cell competition assays

AML cells were transduced to express either GFP or mCherry using the lentiviral pLVX-GFP-P2A-Blasticidin or pLVX-mCherry-P2A-Blasticidin vector, respectively. For sgRNA depletion competition assays, 150,000 GFP+ cells were mixed with 150,000 mCherry+ cells in 12-well plates and spin-infected with lentiCRISPRv2-based lentiviral particles as described above. After 24 h, cells were placed into fresh media in 12-well plates and selected with puromycin for 5 days. Competition assays were conducted for 12 days after selection. Synthetic viability of heme biosynthesis enzymes was validated in a competition format reflecting the setup of the CRISPR screen. GFP+ cells of each genotype (wt or ΔFEM1B) were infected with sgCTRL lentiviral particles and mCherry + cells of each genotype (wt or ΔFEM1B) were infected with sgALAD lentiviral particles. Infected cells were selected with puromycin, after 5 days of selection, cells were counted and mixed 1:1. For example, 150,000 GFP+ wt sgCTRL cells were mixed with 150,000 wt mCherry+ sgALAD and the competitions were conducted for 12 days.

The percentage of mCherry+ cells and GFP+ cells was determined using a BD LSRFortessa instrument, analyzed using FlowJo 10.8.1 and normalized to the sgCNTRL ratio. For drug competition assays 150,000 GFP+ cells were mixed with 150,000 mCherry+ cells in 1 ml of standard culture media in a 12-well plate. Drugs were added during plating and duration of treatment and concentrations are indicated in the corresponding figure legend. The ratio of mCherry+/GFP+ cells was determined using a BD LSRFortessa instrument, analyzed using FlowJo 10.8.1 and normalized to the untreated sample.

### Annexin/PI staining

Induction of apoptosis was followed by determining the fraction of early and late apoptotic cells using Annexin-V/Propidium Iodide staining with the ANNEXIN V-FITC Apoptosis Detection Kit I (BD Pharmingen, 556547) following the manufacturer’s instructions. 1x10^6^ cells were harvested, washed twice in cold 1X PBS and resuspended in 1 ml of binding buffer. 100 µl of this solution was taken for further analysis by adding 3 µl of Annexin-V FITC and 5 µl of PI. Cells were incubated in the dark at room temperature for 15 minutes. 200 µl of binding buffer was added to the samples and samples were immediately measured using a BD LSRFortessa instrument and analyzed using FlowJo 10.8.1. Early apoptotic cells were defined as Annexin-V+/PI- and late apoptotic/necrotic cells were defined as Annexin-V+/PI+.

### Compound screening

To identify synthetic lethal drugs with loss of FEM1B, 20,000 cells (MV4-11 wt or ΔFEM1B) were plated in white 384-well plates in 12.5 µl standard growth media. 1,200 FDA approved compounds (TargetMol Chemicals Inc) were diluted in DMSO and further diluted into 12.5 µl of media and added to the cells to obtain a final concentration of 1 µM drug and 0.5% DMSO using a liquid handler (CyBio Well Vario, Analytik Jena). Cells were incubated with the drugs for 48 hours, after which cell viability was determined using a CellTiter-Glo® Luminescent Cell Viability Assay (Promega, G7570). Cell assay plates and reagents were allowed to equilibrate to room temperature for 30 minutes. CellTiter-Glo reagent was reconstituted according to the manufacturer’s instructions and 25 µl of the solution was added to the 384 well plates using a Multidrop Combi reagent dispenser (Thermo Scientific). Plates were shaken and incubated for 10 minutes, after which the luminescence signal was read on a Perkin Elmer Envision 2104. Data was normalized to control wells in the individual genotype.

### CellTiter-Glo assay

MV4-11 cells (20,000) were seeded in a 384-well plate (total volume 25 μl) with increasing concentrations of Venetoclax (0 nM, 0.75 nM, 1.5 nM, 2.5 nM) and EN106 (0 μM, 0.25 μM, 0.5 μM, 0.75 μM, 1.0 μM, 1.25 μM, 1.5 μM, 1.75 μM, 2.0 μM, 2.5 μM, 3.0 μM) Cell viability was measured after 2 days using a CellTiterGlo Luminescent Cell Viability Assay (Promega) according to manufacturer’s instructions. Luminescence was measured using a plate reader (Tecan, Spark microplate reader). Data were normalized to untreated cells and are represented as mean of triplicates.

### Western blotting

To prevent the aberrant activity of proteases in AML cell lysates, cells were incubated with 2 mM Diisopropyl fluorophosphate (DFP, Thermo Fisher, 115230010) for 15 minutes on ice, followed by two washes in ice cold 1X PBS during harvest. Cell pellets were frozen in liquid N_2_ and stored at -80°C before lysis. For whole cell lysates analysis, cells pellets were lysed in lysis buffer (40mM HEPES pH 7.5, 1% NP40 and 150mM NaCl) supplemented with Roche complete protease inhibitor cocktail (Sigma, 11836145001) and benzonase (EMD Millipore, 70746-4) on ice for 20 min. Lysates were cleared by centrifugation at 21,000 g and samples were then normalized to total protein concentration using Pierce 660 nm Protein Assay reagent (Thermo Fisher, 22660). Next, 2X urea sample buffer (120 mM Tris pH 6.8, 4% SDS, 4 M urea, 20% glycerol and bromophenol blue) was added to the samples. SDS-PAGE and immunoblotting were performed using the indicated antibodies. Images were captured using a ProteinSimple FluorChem M device.

### Small-scale Immunoprecipitations

For small scale co-immunoprecipitations HEK293T cells were transfected, AML cells were lentivirally transduced to express 3X-FLAG tagged proteins using pCS2+ or pLVX-based constructs respectively. Alternatively, CRISPR/Cas9 engineered cell lines with a 3X-FLAG tag at the endogenous BACH1 locus were used. AML cells were harvested using the DFP method as described above to inhibit aberrant protease activity in AML cell lysates. Cell pellets were lysed in IP buffer (40mM HEPES pH 7.5, 0.1% NP40 and 150mM NaCl) supplemented with Roche complete protease inhibitor cocktail (Sigma, 11836145001), PhosSTOP™ (Roche, 4906845001), carfilzomib (Selleckchem, S2853) and benzonase (EMD Millipore, 70746-4) on ice for 20 min. Lysates were cleared by centrifugation at 21,000 g and samples were then normalized to total protein concentration using Pierce 660 nm Protein Assay reagent (Thermo Fisher, 22660). 15-20 µl of ANTI-FLAG M2 Affinity Agarose resin (Sigma-Aldrich, A2220) was added and incubated at 4 °C for 2 h. Lysis, IP and washes were carried out in the presence of hemin as indicated. After four washes of the bound resin with cold IP buffer bound proteins were eluted by addition of 2X urea sample buffer (120 mM Tris pH 6.8, 4% SDS, 4 M urea, 20% glycerol and bromophenol blue) and analyzed by western blotting. Images were captured using a ProteinSimple FluorChem M device.

### Mass spectrometry

For proteomics experiments, lentivirally transduced AML cells expressing 3X-FLAG tagged proteins were scaled up to approximately 1x10^9^ cells and harvested using the DFP method described above. Cell pellets were lysed in 3X pellet volume of IP buffer and the IP was carried out as described above with 50µl of anti-FLAG resin. After three washes with IP buffer, three additional washes with PBS were performed. All of the remaining liquid was removed with a crushed gel tip and the beads were flash-frozen in liquid N_2_ and sent to further processing at UC San Diego Proteomics Facility. Protein samples were diluted in TNE (50mM Tris pH8.0, 100mM NaCl, 1mM EDTA) buffer. RapiGest SF reagent (Waters) was added to the mix to a final concentration of 0.1%, and the samples were boiled for 5 min. TCEP was added to 1 mM (final concentration) and the samples were incubated at 37 °C for 30 min. Subsequently, the samples were carboxymethylated with 0.5 mg ml−1 of iodoacetamide for 30 min at 37 °C followed by neutralization with 2 Mm TCEP (final concentration). The proteins samples were then digested with trypsin (trypsin:protein ratio, 1:50) overnight at 37 °C. RapiGest was degraded and removed by treating the samples with 250 mM HCl at 37 °C for 1 h followed by centrifugation at 14,000 rpm for 30 min at 4 °C. The soluble fraction was then added to a new tube, and the peptides were extracted and desalted using C18 desalting columns (Thermo Fisher Scientific, PI-87782). Peptides were quantified using BCA assay and a total of 1 μg of peptides were injected for LC–MS analysis. Trypsin-digested peptides were analysed by ultra-high-pressure liquid chromatography (UPLC) coupled with tandem mass spectroscopy (LC–MS/MS) using nano-spray ionization. The nanospray ionization experiments were performed using a TimsTOF 2 pro hybrid mass spectrometer (Bruker) interfaced with nanoscale reversed-phase UPLC (EVOSEP ONE).

The Evosep method of 30 samples per day was performed using a 10 cm × 150 μm reversed-phase column packed with 1.5 μm C18-beads (PepSep, Bruker) at 58 °C. The analytical columns were connected with a fused silica ID emitter (10 μm inner diameter, Bruker Daltonics) inside a nanoelectrospray ion source (captive spray source, Bruker). The mobile phases comprised 0.1% formic acid as solution A and 0.1% formic acid/99.9% acetonitrile as solution B. The MS settings for the TimsTOF Pro 2 were as follows: the DIA-PASEF method for proteomics. The values for mobility-dependent collision energy were set to 10 eV. No merging of TIMS scans was performed. The ion mobility (IM) was set between 0.85 (1/k0) and 1.3 (1/k0) with a ramp time of 100 ms. Each method includes one IM window per DIA-PASEF scan with variable isolation window at 20 amu segments; 34 PASEF MS/MS scans were triggered per cycle (1.38 s) with a maximum of 7 precursors per mobilogram. Precursor ions in an m/z range of between 100 and 1,700 with charge states ≥3+ and ≤8+ were selected for fragmentation. Protein identification and label-free quantification were performed using Spectronaut 18.0 (Biognosys). Data analysis: For proteomics experiments presented in Figure 2 and Figure S2 results were filtered by subtraction of an empty vector control sample after DIA analysis. NaN values were replaced by 0. Next, contaminant proteins were excluded from the analysis if they appeared in ≥25% of datasets in the CRAPOME database ^95^. Detected intensities were normalized to bait, log_2_-transformed and plotted. For the proteomics experiment assessing heme dependent interactions of BACH1 presented in Figure 4 samples were run as technical triplicates on the mass spectrometer. Data was processed as described above. Values relative to the average DIA quantity of untreated samples are displayed as a heat map for a subset of known and validated BACH1 interactors as well as the CUL2^FEM1B^ complex. Analysis of MS-data comparing the interactome of BACH1^wt^ and BACH1^C646S^ was performed using default statistical testing of technical MS replicates using Spectronaut (Biognosys) without initial filtering of an EV control sample or removal of contaminant proteins.

### Heme/Iron measurements

Expi293F cells were transfected with FEM1B and BACH1 overexpression constructs as indicated. Cell pellets were lysed and anti-FLAG IP was performed as described above in the presence of 20µM hemin. After two washes with hemin containing buffer, beads were washed three times in TBS (150mM NaCl, 40mM Tris-HCl pH 7.5) to remove excess hemin and proteins were then eluted using 3XFLAG-peptide (MedChemExpress, HY-P0319). Heme measurements were then performed by using an apo-Peroxidase (apoHRP) based assay ^96^. Briefly, the commercial apoHRP (Calzyme, 239A0000) was extracted by acid acetone solution in order to eliminate any possible residual HRP activity. 5µl of eluate were then added to 10µl of 50µM apoHRP and 85µl 1XPBS. After a 10 minute incubation on ice, 10µl of this reaction were transferred in triplicates to a clear 96-well plate and 200µl of TMB substrate (Vector laboratories, SK-4400) was added. The plate was protected from light and incubated for 15 minutes at room temperature, followed by measurement of absorbance at 650nm. A standard curve was determined using defined concentrations of hemin incubated with apoHRP.

For detection of iron by inductively coupled plasma spectroscopy (ICP), eluted proteins were digested overnight with Chymotrypsin (Promega, V1061) and denatured and precipitated the next day by adding HNO_3_ to a final concentration of 3%, followed by centrifugation at 21000g for 10 minutes. Inductively coupled plasma spectroscopy was performed using the supernatants on a Perkin Elmer 5300 DV. Standard curves (0, 0.01, 0.1, and 1 mg/mL) were prepared for several transition metals (Sigma, 04330-100ML), and metal concentrations were determined using a linear fit from the standard curves.

### Protein expression and purification

<u>FEM1B-ElonginB/C</u>: 6xHis-MBP-TEV-FEM1B and Elongin B and C were co-expressed in LOBSTR *E. coli* cells grown in LB broth medium. Bacterial cells were grown at 37 °C to an OD_600_ of 0.6, induced with 0.4 mM isopropyl-β-D-1-thiogalactopyranoside and proteins were expressed at 16 °C overnight. The cells were harvested (7808 x g) and lysed by sonication in buffer (50 mM Tris, pH 8.0, 250 mM NaCl, 10 mM imidazole, 2 mM MgSO_4_, 5 mM β-mercaptoethanol) supplemented with benzonase nuclease, EDTA-free cOmplete protease inhibitor cocktail and lysozyme. After centrifugation (36000 x g, 1h), the cleared cell lysate was added to Ni-NTA resin, washed with lysis buffer containing 2 M NaCl, and then with lysis buffer containing 20 mM imidazole. The protein complex was eluted with 250 mM imidazole and subsequently purified by ion-exchange chromatography (20 mM Tris, pH 8.0, gradient from 0.02 M to 1 M NaCl in 10 column volumes) on a HiTrap Q HP anion exchange column (Cytiva, 17115401) followed by size-exclusion chromatography (20 mM HEPES, pH 7.5, 150 mM NaCl, 2 mM DTT) on a HiLoad 16/600 Superdex 200 pg column (Cytiva, 28989335). FEM1B mutants (K16E and K16E/L18A/L25A) were purified as described above.

<u>BACH1 CT</u>: BACH1 CT-6xHis was expressed in LOBSTR *E. coli* cells. The bacterial cells were grown in LB broth medium at 37 °C to an OD_600_ of 0.6, induced with 0.4 mM isopropyl-β-D-1-thiogalactopyranoside and proteins were expressed at 16 °C overnight. The cells were harvested (7808 x g) and lysed by sonication in buffer (50 mM Tris, pH 8.0, 250 mM NaCl, 10 mM imidazole, 2 mM MgSO_4_, 5 mM β-mercaptoethanol) supplemented with benzonase nuclease, EDTA-free cOmplete protease inhibitor cocktail and lysozyme. After centrifugation (36000 x g, 1h), the cleared cell lysate was added to Ni-NTA resin, washed with lysis buffer containing 2 M NaCl, warm lysis buffer containing 5 mM ATP and 10 mM MgCl_2_ and then with lysis buffer containing 20 mM imidazole. The protein complex was eluted with 250 mM imidazole and subsequently purified by ion-exchange chromatography (20 mM Tris, pH 8.0, gradient from 0.02 M to 1 M NaCl in 10 column volumes) on a HiTrap Q HP anion exchange column (Cytiva, 17115401) followed by size-exclusion chromatography (20 mM HEPES, pH 7.5, 150 mM NaCl, 2 mM DTT) on a HiLoad 16/60 Superdex 75 prep grade column (Cytiva, 17106801).

<u>CUL2-Rbx1</u>: 6xHis-mysB-TEV-StrepII-CUL2 (Δ117-134) was co-expressed with 6xHis-TEV-Rbx1 in LOBSTR *E. coli* cells grown in LB broth medium. Bacterial cells were grown at 37 °C to an OD_600_ of 0.6, induced with 0.4 mM isopropyl-β-D-1-thiogalactopyranoside and proteins were expressed at 16 °C overnight. The cells were harvested (7808 x g) and lysed by sonication in 50 mM Tris, pH 8.0, 250 mM NaCl, 20 mM imidazole, 2 mM MgSO_4_, 5 mM β-mercaptoethanol, benzonase nuclease, EDTA-free cOmplete protease inhibitor cocktail and lysozyme. After centrifugation (36000 x g, 1h), the cleared cell lysate was added to Ni-NTA resin, washed with lysis buffer containing 2 M NaCl, and then with lysis buffer containing 20 mM imidazole. The protein complex was eluted with 250 mM imidazole, and the 6xHis-tag was cleaved at 4 °C overnight (TEV produced in-house, UC Berkeley MacroLab; ∼1:50 w/w). The solution was incubated with StrepTactin 4Flow XT resin (IBA, 2-5010) for 1 hour in batch on a nutator in 50 mL conical tubes. Beads were loaded onto a gravity purification column (BioRad, 7374011), washed once with ∼5-column volumes of buffer (50 mM HEPES, pH 7.5, 250 mM NaCl, 2 mM DTT), and the protein was eluted in buffer containing 50 mM biotin. The eluate was concentrated and further purified by size-exclusion chromatography (20 mM HEPES, pH 7.5, 250 mM NaCl, 2 mM DTT) on a Superdex 200 Increase 10/300 GL column (Cytiva, 28990944).

<u>CUL2-Rbx1-FEM1B-ElonginB/C complex for *in vitro* ubiquitylation assays:</u> The purified 6xHis-MBP-TEV-FEM1B-Elongin B/C complex was mixed with purified CUL2-Rbx1 in a 1:1 ratio, and the 6xHis-Tag was cleaved with TEV protease (produced in-house, UC Berkeley MacroLab; ∼1:50 w/w) at 4 °C overnight. Subsequently, the complex was purified by size-exclusion chromatography (20 mM HEPES, pH 7.5, 150 mM NaCl, 2 mM DTT) on a Superose 6 10/300 GL column (Cytiva, 29091596). The fractions were pooled, flash-frozen, and stored at -80 °C for future use.

<u>CUL2-Rbx1-FEM1B-ElonginB/C-BACH1 CT complex for structure determination:</u> The purified 6xHis-MBP-TEV-FEM1B-Elongin B/C complex and BACH1 CT were mixed in a 1:1 ratio in the presence of 10 μM Hemin, and the 6xHis-Tag was cleaved with TEV protease (produced in-house, UC Berkeley MacroLab; ∼1:50 w/w) at 4 °C overnight. Subsequently, the complex was purified by size-exclusion chromatography (20 mM HEPES, pH 7.5, 150 mM NaCl, 10 μM Hemin, 2 mM DTT) on a Superdex 200 Increase (10/300) column (Cytiva, 28990944). The purified FEM1B-ElonginB/C-BACH1 CT complex was incubated with CUL2-Rbx1 for 2 hours (20 mM HEPES, pH 7.5, 150 mM NaCl, 10 μM Hemin, 2 mM DTT) and purified via a Superose 6 10/300 column (Cytiva, 29091596) in cryo-EM buffer (20 mM HEPES, pH 7.5, 150 mM NaCl, 10 μM Hemin, 2 mM DTT). The Fractions were pooled, concentrated to 7.5 mg/mL and immediately used for Cryo-EM grid preparation.

### *In vitro* ubiquitylation assays

Recombinant CUL2-Rbx1-FEM1B-ElonginB/C complex was first subjected to a neddylation reaction. For each neddylation reaction (50 μL reaction volume), 5 μM recombinant CUL2-Rbx1-FEM1B-ElonginB/C was incubated with 25 mM Nedd8, 0.5 μM UBA3, 1 μM UBE2M, 0.2 mM DTT and 1 x energy mix (150 mM creatine phosphate, 20 mM ATP, 20 mM MgCl_2_, 2 mM EGTA, pH to 7.5 with KOH) in UBA buffer (25 mM Tris-HCl, pH 7.5, 50 mM NaCl, 10 mM MgCl_2_) for 30 minutes at 30 °C. *In vitro* ubiquitylation assays were performed in 10 μl reaction volume. 1 μM neddylated CUL2-Rbx1-FEM1B-ElonginB/C was incubated with 0.3 μM BACH1 CT, 0.25 μM E1, 2.5 μM UBE2R1, 1 mg/mL Ubiquitin, 1 mM DTT and 1.5 μl of energy mix (150 mM creatine phosphate, 20 mM ATP, 20 mM MgCl_2_, 2 mM EGTA, pH to 7.5 with KOH) in UBA buffer (25 mM Tris-HCl, pH 7.5, 50 mM NaCl, 10 mM MgCl_2_) for 1 hour at 30 °C. *In vitro* ubiquitylation assays were performed at different hemin concentrations as indicated. Reactions were quenched in 2X Urea sample buffer (120 mM Tris, pH 6.8, 4% SDS, 20% glycerol, and bromophenol blue) and resolved 12% SDS-acrylamide gels or SDS-acrylamide 4-20% gradient gels. After immunoblotting, images were captured using the ProteinSimple FluorChem M device.

### Biolayer Interferometry

6x-His-MBP-FEM1B or BACH1 CT-6xHis, respectively, were biotinylated with NHS-Biotin (EZ-Link NHS-Biotin, ThermoFisher) in a 1:1 ratio (20 mM HEPES, pH 7.5, 150 mM NaCl, 2 mM TCEP) for 2 hours on ice. Excess Biotin was removed by desalting (Zeba Spin desalting columns, 7K MWCO, 0.5 mL, ThermoFisher). Binding interactions were analyzed on an Octet RED 384 instrument in 384-well plates (greiner, 781900) at 27 °C 50 μl volume. Octet Streptavidin Biosensors (SARTORIUS, 18-5019) were hydrated in buffer (20 mM HEPES, pH 7.5, 150 mM NaCl, 2 mM DTT, 0.1% BSA, 0.02% Tween20) for 10 minutes before use. Biotinylated ligand was diluted in the respective buffer and immobilized onto Streptavidin Biosensors to a loading response ∼ 1.0 - 1.5 nm. Following a wash step in the respective buffer, a baseline was established (60 s). Association measurements (240 s) were performed by transferring loaded biosensors into wells with the respective analyte. Depending on the experiment, either increasing protein concentrations (serial dilution) at a constant hemin concentration in the buffer or increasing hemin concentrations at constant protein concentration was measured. Dissociation was monitored by transferring biosensors to buffer (240 s). Data analysis was performed using Octet DataAnalysis software (v11.1.0.4). Reference subtraction was performed using a parallel sensor in buffer to correct for signal drift, the curves were aligned to the baseline, and interstep correction was applied prior to fitting. Data collected across multiple protein or hemin concentrations were fitted using a 2:1 heterogeneous ligand-binding model. The apparent K_D_ was determined by steady-state analysis by plotting equilibrium response against analyte concentration and fitting to a binding isotherm. Similar results in n=2 independent experiments.

### Cryo-EM sample preparation, data collection, and processing

Cryo-EM samples were mixed with a final concentration of 0.02% (w/v) fluorinated octylmaltoside (Anatrace, O310F) immediately before cryo-freezing to prevent protein denaturation at the air–water interface. 2.6 μL of the sample was subsequently applied to a glow-discharged 300-mesh Quantifoil R1.2/1.3 grid and incubated for 15 s before being blotted and plunge-vitrified in liquid ethane cool-protected by liquid nitrogen. To reduce particle aggregation and denaturation, the CUL2^FEM1B^-BACH1^CT^ complex sample was cryo-preserved on Quantifoil R1.2/1.3 grids coated with 2 nm amorphous carbon film (Electron Microscopy Sciences). Grid freezing was performed using a Mark IV Vitrobot (Thermo Fisher Scientific) system operating at 12 °C and 100% humidity. Cryo-EM data were collected using 300 kV Titan Krios G3 microscopes (Thermo Fisher Scientific) equipped with a BIO Quantum energy filter (slit width 20 eV). Data were collected using SerialEM software at a nominal magnification of 105,000x with a pixel size of 0.42 Å/pixel or 0.414 Å/pixel. Movies were recorded using a 6k x 4k Gatan K3 Direct Electron Detector operating in super-resolution CDS mode. Each movie was composed of 40 subframes with a total dose of 50 e^-^/A^2^, resulting in a dose rate of ∼1.25 e^-^/A^2^*sec. Data processing, including motion correction, CTF estimation, particle picking, 2D class averaging, and 3D refinement, was performed using cryoSPARC v.5.0 workflow, using mostly default settings. All movies were 2x binned and patch motion corrected. After particle picking and several iterations of 2D class averaging, the initial 3D volume was calculated using several rounds of *ab initio* 3D reconstruction and classification. For local refinement of the FEM1B-BACH1 interface, density regions corresponding to BACH1 and the N-terminal 377 residues of FEM1B were isolated from the 3.8Å monomeric density map, density for the remaining regions was subtracted from particle images using the Particle Subtraction program (cryoSPARC v5.0). Local refinement (cryoSPARC v5.0) was subsequently performed and a 5.9Å map was generated. AF3 model of FEM1B-BACH1 complex was then fitted into the local refinement map. Default B-factor sharpening in cryoSPARC v5.0 was used to generate the final deposited maps. A full workflow of cryo-EM data processing is outlined in **Figure S4**.

### Model building and structural analysis

The atomic models of the dimeric and monomeric CUL2FEM1B-BACH1CT complex was built based on the published structure of the CUL2FEM1B complex (PDB 8WQB). The structural model of the FEM1B-BACH1 complex was predicted using AlphaFold3 ^97^ with the full-length FEM1B, two copies of BACH1 (residues 611-653), and heme. The predicted AF3 model was fitted into the FEM1B-BACH1 local refinement map. Key residues located at the protein-protein interface were identified and subsequently validated using in vitro biochemical and cell-based assays.

### Multiple sequence alignments (MSAs)

MSA’s were performed using Clustal Omega ^98^. The FEM1B conservation score was determined by aligning FEM1B human, mouse, rat, bovine, chicken, and frog orthologs. UniProt sequence identifiers are as follows: (species, FEM1 protein, FEM1 accession): human (Homo sapiens, FEM1A, Q9BSK); human (Homo sapiens, FEM1C, Q96JP0); human (Homo sapiens, FEM1B, Q9UK73); mouse (Mus musculus, FEM1B, Q9Z2G0); rat (Rattus norvegicus, FEM1B, P0C6P7); bovine (Bos taurus, FEM1B, A0ABI0NTK6); chicken (Gallus gallus, FEM1B, Q5ZM55); frog (Xenopus laevis, FEM1B, Q6GPE5).

### MBP pulldown assays

Binding experiments were done at room temperature in 200µl of buffer (40mM HEPES pH 7.5, 150mM NaCl, 0.1% NP40 substitute, 1mM of DTT) with 1µM final concentration of MBP-^6XHis^FEM1B and 10µM BACH1^CT^. Hemin was added to binding buffer as indicated. Binding reactions were added to 15 µl of amylose resin and left to bind for 60 minutes at room temperature. Beads were washed 3x in 500µl binding buffer and proteins eluted in 2x urea sample buffer and analyzed by SDS-PAGE and western blotting as indicated. All inputs are of total protein.

### RNAseq sample preparation and analysis

MV4-11 cells were depleted of FEM1B or BACH1 by lentiviral transduction of pLKO.1 shRNA plasmids (Addgene, 8453). 5x10^5^ cells were spinfected with such lentiviral particles as described above with a media change the following day. Cells were harvested and flash frozen after 72 h. Cell pellets were resuspended in DNA/RNA Shield (Zymo Research, R1100-50) and sent to Plasmidsaurus, which utilizes Illumina sequencing and a 3’ end counting approach. Quality of the fastq files was assessed using FastQC v0.12.1. Reads were then quality filtered using fastp v0.24.0 with poly-X tail trimming, 3’ quality-based tail trimming, a minimum Phred quality score of 15, and a minimum length requirement of 50 bp. Quality-filtered reads were aligned to the reference genome using STAR aligner v2.7.11 with non-canonical splice junction removal and output of unmapped reads, followed by coordinate sorting using samtools v1.22.1. PCR and optical duplicates were removed using UMI-based deduplication with UMIcollapse v1.1.0. Alignment quality metrics, strand specificity, and read distribution across genomic features were assessed using RSeQC v5.0.4 and Qualimap v2.3. Gene-level expression quantification was performed using featureCounts (subread package v2.1.1) with strand-specific counting, multi-mapping read fractional assignment, exons and three prime UTR as the feature identifiers, and grouped by gene_id. Differential expression analysis was done with edgeR v4.0.16 using standard practice including filtering for low-expressed genes with edgeR::filterByExpr with default values.

### qPCR

MV4-11 cells were depleted of the indicated genes by lentiviral transduction of pLKO.1 shRNA plasmids (Addgene, 8453). 5x10^5^ cells were spinfected with such lentiviral particles as described above with a media change the following day. Cells were harvested and flash frozen after 72 h. Total RNA was purified using a nucleospin RNA kit (Macherey-Nagel, 740955) following the manufacturers instructions. cDNA was generated using a RevertAid First Strand cDNA Synthesis kit (Thermo Fisher Scientific, K1622) and RT-qPCRs were performed using 2×KAPA SYBR Fast qPCR master mix (Roche, KK4602) on a LightCycler 480 II Instrument (Roche) using. Fold changes in expression were calculated using the ΔΔCt method. Primers used for qPCR are listed in Supplementary Table 1.

### ChIP-qPCR

1.5x10^7^ MV4-11 cells were crosslinked per IP in 20ml of media containing 1% paraformaldehyde (ThermoFisher, 28908) for 5 min at room temperature and quenched with 125 mM glycine for 5 min. DFP treatment (as described above) was carried out in parallel to inhibit aberrant protease activity in AML lysates. All buffers contained Roche complete protease inhibitor cocktail (Sigma, 11836145001). Cells were washed twice in cold 1x PBS and immediately resuspended in 1 ml of lysis buffer (50 mM HEPES pH 7.5, 140 mM NaCl, 1 mM EDTA, 10% glycerol, 0.5% NP40, 0.25% Triton X-100) and incubated for 10 minutes on a rotator at 4°C. Nuclei were collected by centrifuging at 1700 g for 5 min, followed by a 10 minute incubation in 1 ml of wash buffer (10 mM TrisHCl pH 8.1, 100 mM NaCl, 1 mM EDTA, pH 8.0).

Nuclei were again collected by centrifugation, pellet gently rinsed and later resuspended with 1 ml of shearing buffer (50 mM Tris Cl, pH 7.5, 10 mM EDTA, 0.1% SDS). Samples were transferred to AFA tube with fiber (Covaris # 520130) and sonicated in a Covaris S220 focused ultrasonicator using default settings (140 power, 5% duty cycle, 200 bursts/cycle, 5 °C waterbath temperature). Sheared chromatin was cleared by centrifugation at 21,000 g for 10 minutes, followed by pre-clear with 25 µl of Protein G Dynabeads™ (ThermoFisher, 10004D) for 2 hr at 4°C. DNA quantification was performed using the Qubit broad range assay kit (Thermo Fisher, Q33265). 5 µg of pre-cleared chromatin was diluted to 500 µl volume with shearing buffer and supplemented to a final concentration of 150 mM NaCl and 1% Triton X 100. 5 µl of this sample was taken as input and added to 95µl of 150 mM NaCl buffer with 0.5 µl of Proteinase K (ThermoFisher, EO0491) and 1 µl of RNAse A (Thermo Fisher, EN0531). 0.65 µg of rabbit monoclonal anti-BACH1 (E4E7B, Cell Signaling Technology #33059) or normal rabbit IgG (Cell Signaling Technology #2729) were added to the IP samples and incubated overnight on a rotator at 4 °C. In parallel, 25 µl of Protein G Dynabeads™ per sample were blocked in BSA. Next day, blocked beads were washed three times, added to IP samples and incubated for 2 h at 4 °C. The following washes were carried out for 10 minutes rotating at 4°C and using a DynaMag™ magnetic rack (Thermo Fisher, 12321D). Beads were washed in 1 ml of low salt wash buffer (20 mM Tris 8.0, 150 mM NaCl, 2 mM EDTA, 1% Triton X-100 and 0.1% SDS), followed by 1 ml of high salt wash buffer (20 mM Tris 8.0, 500 mM NaCl, 2 mM EDTA, 1% Triton X-100 and 0.1% SDS), followed by 1 ml of LiCl buffer (20 mM Tris 8.0, 250 mM LiCl, 1 mM EDTA, 1% deoxycholate and 1% Nonidet P-40), and one final wash 1× TE. Samples were then eluted by addition of 100 µl of 150mM NaCl with 0.5 µl proteinase K and 1 µl RNAse A per sample. Samples were reverse crosslinked overnight at 65 °C. DNA was then purified and concentrated using the Zymo Oligo Clean & Concentrator Kit (Zymo Research, D4060) and eluted in 50 µl of nuclease-free H_2_O. 2 µl of DNA were used for qPCR, which was performed as described above. Primers used for qPCR can be found in Supplementary Table 1.

### Quantification and statistical analysis

All quantifications are presented as mean ± standard deviation or standard error of means as indicated in the figure legends. Significance was determined by two-tailed t tests or one-sample t-tests in Graphpad Prism. ns p > 0.05, *p ≤ 0.05; **p ≤ 0.01; ***p ≤ 0.001, ****p ≤ 0.0001. The number of biological replicates is specified in the corresponding figure legends.

## Key resources table

| REAGENT or RESOURCE | SOURCE | IDENTIFIER |
| --- | --- | --- |
| **Antibodies** |  |  |
| rabbit monoclonal DYKDDDDK Tag Antibody | Cell Signaling Technology | Cat# 2368; RRID:AB_2217020 |
| rabbit monoclonal anti-GAPDH (14C10) | Cell Signaling Technology | Cat# 2118S; RRID:AB_561053 |
| mouse monoclonal anti-FLAG clone M2 | Sigma-Aldrich | Cat# F1804; RRID:AB_262044 |
| rabbit polyclonal anti-CUL2 | Bethyl | Cat# A302-476A; RRID:AB_1944215 |
| rabbit monoclonal anti-FNIP1 | Abcam | Cat# ab134969; RRID:AB_3170229 |
| rabbit polyclonal anti-FEM1B | Proteintech | Cat# 11030-1-AP; RRID:AB_2102637 |
| rabbit polyclonal anti-BACH1 | Proteintech | Cat# 14018-1-AP; RRID:AB_2274498 |
| mouse monoclonal anti-BACH1 | Santa Cruz | Cat# sc-271211; RRID:AB_10608972 |
| normal rabbit IgG | Cell Signaling Technology | Cat# #2719; RRID:AB_1031062 |
| rabbit polyclonal anti-FBXO22 | Proteintech | Cat# 13606-1-AP; RRID:AB_2104403 |
| mouse monoclonal anti-COX4 | Abcam | Cat# ab14744; RRID:AB_301443 |
| mouse monoclonal anti-Alpha Tubulin | Sigma-Aldrich | Cat# CP06; RRID:AB_2617116 |
| goat anti-rabbit IgG (H+L) HRP | Vector Laboratories | Cat# PI-1000-1; RRID:AB_2916034 |
| sheep anti-mouse IgG (H+L) HRP | Sigma-Aldrich | Cat# A5906; RRID:AB_258264 |
| goat anti-mouse IgG light-chain-specific HRP conjugated | Jackson ImmunoResearch | Cat# 115-035-174; RRID:AB_2338512 |
| APC anti-rat CD90/mouse CD90.1 (Thy-1.1) | Biolegend | Cat# 202526; RRID:AB_1595470 |
| **Bacterial and virus strains** |  |  |
| E.coli: One Shot Stbl3 Chemically competent cells | Thermo Fisher Scientific | Cat#C737303 |
| E.coli LOBSTR-BL21(DE3)-RIL | Vector Laboratories | NC1789768 |
| **Biological samples** |  |  |
| Bone marrow aspirates of 675 AML patients and 64 healthy control bone marrow mononuclear cells | This study | N/A |
| **Chemicals, peptides, and recombinant proteins** |  |  |
| Recombinant Human E1/UBA1 Protein | Laboratory of Michael Rapé | N/A |
| Recombinant Human APPBP1/UBA3 Complex (NEDD8 E1) Protein, CF | R&D Systems | Cat# E-313 |
| Recombinant Human UBE2R1 | Laboratory of Michael Rapé | N/A |
| Recombinant Human NEDD8 | Laboratory of Michael Rapé | N/A |
| Recombinant Human UBE2R1 | This paper | N/A |
| Recombinant Human CUL2 (CUL2<sup>Δ117-134</sup>-RBX1) | This paper | N/A |
| Recombinant Human His-MBP | Laboratory of Michael Rapé | N/A |
| Recombinant TOM20 (62-127) | <sup>36</sup> | N/A |
| Recombinant Human 6XHis-MBP-FEM1B (wt, K16E, L18A/L25A, K16E/L18A/L25A) | This paper | N/A |
| Recombinant Human BACH1-CT-6XHis (506-684) (wt, L617A/L620A, C646S) | This paper | N/A |
| Recombinant Human Ubiquitin Protein, CF | R&D Systems | Cat# U-100H |
| apoPeroxidase | Calzyme | Cat# 239A0000 |
| Chymotrypsin | Promega | Cat# V1061 |
| Transition metal mix 1 for ICP | Sigma | Cat# 04330-100ML |
| TMB Substrate Kit, Peroxidase (HRP), (3,3', 5,5'-tetramethylbenzidine) | Vector laboratories | Cat# SK-4400 |
| Deferiprone | Selleckchem | Cat# S4067 |
| Hemin | Sigma-Aldrich | Cat# H9039 |
| Fe(III) Protoporphyrin IX dimethyl ester chloride | Frontier Specialty Chemicals | Cat# P40311 |
| Fe(III) Deuteroporphyrin IX chloride | Frontier Specialty Chemicals | Cat# D40654 |
| Cobaltic(III) Protoporphyrin IX Chloride | Cayman Chemicals | Cat# 33794 |
| Mn(III) Protoporphyrin IX Chloride | Cayman Chemicals | Cat# 30808 |
| Venetoclax (ABT-199) | Adooq Bioscience | Cat# A12500 |
| EN106 | MedChemExpress | Cat# HY-W229874 |
| EZ-Link NHS-Biotin | Thermo Fisher | Cat# 20217 |
| Creatine phosphate | Sigma-Aldrich | Cat#10621714001-5G |
| cOmplete™, EDTA-free protease inhibitor cocktail tablets from Roche | Sigma-Aldrich | Cat#11873580001 |
| Roche PhosSTOP™ | Sigma-Aldrich | Cat#4906845001 |
| Polyethylenimine (PEI), Linear, MW 25000, Transfection Grade | Polysciences | Cat#23966-1 |
| Carfilzomib (PR-171) | Selleck Chemicals | Cat#S2853 |
| TEV protease | UCB QB3 MacroLab | N/A |
| Recombinant Cas9 Protein | UCB QB3 MacroLab | N/A |
| Alt-R™ HDR Enhancer V2 | IDT Technologies | Cat# 10007910 |
| MLN4924 | Selleck Chemicals | Cat# NC1533706 |
| Benzonase | EMD Millipore | Cat# 70746-4 |
| Pierce™ 16% Formaldehyde (w/v), Methanol-free | ThermoFisher | Cat# 28908 |
| AmpliTaq Gold™ DNA Polymerase | Thermo Fisher | Cat# 4311820 |
| Bafilomycin A1 (Baf-A1) | SelleckChem | Cat# S1413 |
| ABT-263 | AdooQ Bioscience | Cat# A10022 |
| ABT-737 | AdooQ Bioscience | Cat# A10255 |
| S63845 | SelleckChem | Cat# S8383 |
| S55746 | SelleckChem | Cat# S8759 |
| Z-VAD-FMK | AdooQ Biosciences | Cat# A12373 |
| Proteinase K, recombinant, PCR grade | ThermoFisher | Cat#EO0491 |
| **Critical commercial assays** |  |  |
| Pierce 660nm Protein Assay Reagent | Thermo Fisher | Cat# 22660 |
| Ionic Detergent Compatibility Reagent | Thermo Fisher | Cat# 22663 |
| KAPA SYBR FAST Uni | Roche Diagnostics Corporation | Cat# 07959397001 |
| NucleoSpin RNA, Mini kit for RNA purification | MACHEREY-NAGEL | Cat# 740955.250 |
| Qubit dsDNA High Sensitivity assay kit | Thermo Fisher | Cat# Q33230 |
| Qubit dsDNA Broad Range assay kit | Thermo Fisher | Cat# Q33265 |
| CellTiter-Glo® Luminescent Cell Viability Assay | Promega | Cat# G7571 |
| Octet® Streptavidin (SA) Biosensor | Sartorius | Cat# 18-5019 |
| BD Pharmingen™ FITC Annexin V Apoptosis Detection Kit I | BD Biosciences | Cat# 556547 |
| QIAamp DNA Blood Maxi Kit | QIAGEN | Cat# 51194 |
| AMPure XP Beads for DNA Cleanup | Beckman Coulter | Cat# A63881 |
| **Deposited data** |  |  |
| RNA-seq MV4-11 shCTRL, shFEM1B, shBACH1 | This study | GSE328744 |
| Proteomics datasets | This study | PXD077617 |
| Cryo-EM map of CUL2<sup>FEM1B</sup>-BACH1<sup>CT</sup> (dimer) | This study | EMD-76876 |
| Cryo-EM map and model of CUL2<sup>FEM1B</sup>-BACH1<sup>CT</sup> (monomer) | This study | PDB ID 12ZS; EMD-76897 |
| Cryo-EM map and model for local refinement of the FEM1B-BACH1 interaction modules | This study | PDB 12ZX, EMD-76910 |
| **Experimental models: Cell lines** |  |  |
| HEK293T | UCB Tissue Culture Facility | RRID:CVCL_0063 |
| HEK293T-BACH1-3xFLAG | This paper | N/A |
| HEK293T-ΔFEM1B | <sup>37</sup> | N/A |
| HEK293T-ΔBACH1 | This paper | N/A |
| HEK293T-ΔFBXL17 | <sup>59</sup> | N/A |
| Expi293F | UCB Tissue Culture Facility | RRID:CVCL_D615 |
| THP1 | UCB Tissue Culture Facility | RRID:CVCL_0006 |
| THP1-ΔFEM1B | This paper | N/A |
| THP1-BACH1-3XFLAG | This paper | N/A |
| THP1-Cas9 | This paper | N/A |
| THP1-ΔFEM1B-Cas9 | This paper | N/A |
| MV4-11 | UCB Tissue Culture Facility | RRID:CVCL_0064 |
| MV4-11-ΔFEM1B | This paper | N/A |
| MV4-11-BACH1-3XFLAG | This paper | N/A |
| MV4-11-Cas9 | This paper | N/A |
| MV4-11-ΔFEM1B-Cas9 | This paper | N/A |
| MV4-11-BACH1<sup>C646S</sup> | This paper | N/A |
| MV4-11-BACH1<sup>L617A/L620A</sup> | This paper | N/A |
| MOLM13 | UCB Tissue Culture Facility | RRID:CVCL_2119 |
| MOLM13-ΔFEM1B | This paper | N/A |
| MOLM13-BACH1<sup>C646S</sup> | This paper | N/A |
| MOLM13-BACH1<sup>L617A/L620A</sup> | This paper | N/A |
| **Experimental models: Organisms/strains** |  |  |
| N/A |  |  |
| **Oligonucleotides** |  |  |
| See Supplementary Tables 1-3 for a list of oligonucleotides |  |  |
| **Recombinant DNA** |  |  |
| pCS2-BACH1<sup>3xFLAG</sup> wt | This paper | N/A |
| pCS2-BACH1<sup>3xFLAG</sup> (506-684) | This paper | N/A |
| pCS2-BACH1<sup>3xFLAG</sup> (Δ506-684) | This paper | N/A |
| pCS2-BACH1<sup>3xFLAG</sup> (C646S, C621S/C625S, L617A, L620A, L617A/L620A, L617A/L620A/V624A, Y641A) | This paper | N/A |
| pCS2-<sup>3xFLAG</sup>FEM1A wt | 35 | N/A |
| pCS2-FBXL17<sup>3xFLAG</sup> wt | 59 | N/A |
| pCS2-FBXO22<sup>3xFLAG</sup> wt | This paper | N/A |
| pCS2-ALAS1<sup>3xFLAG</sup> wt | This paper | N/A |
| pCS2-ALAD<sup>3xFLAG</sup> wt | This paper | N/A |
| pCS2-TOM20<sup>HA</sup> wt | 36 | N/A |
| pCS2-cytoTOM20<sup>HA</sup> (25-145) | 36 | N/A |
| pCS2-<sup>3xFLAG</sup>FEM1B wt | This paper | N/A |
| pCS2-<sup>3xFLAG</sup>FEM1B (L18A/L25A, K16E, K16E/L18A/L25A, T19E, A22E, K60A, W93A, E102A, F130A, R126Q, C186S, V391A/Q394A, F501A/H502A, L597A, R126A/L597A) | This paper | N/A |
| pCS2-<sup>3xFLAG</sup>FEM1B (F549D/V584D/I587D/L588D) | 60 | N/A |
| pCS2-AKAP1(1-30)-(GGGS)<sub>3</sub>-FEM1B wt | This paper | N/A |
| pCS2-AKAP1(1-30)-(GGGS)<sub>6</sub>-FEM1B wt | This paper | N/A |
| PLVX-EF1α-<sup>3xFLAG</sup>FEM1B (wt, R126A/L597A) | This paper | N/A |
| PLVX-EF1α-BACH1<sup>3xFLAG</sup> (wt, C646S) | This paper | N/A |
| PLVX-Tet-one-BACH1-GFP-IRES-mCherry | This paper | N/A |
| PLVX-Tet-3G-BACH1-mNeonGreen-IRES-mScarlet3 | This paper | N/A |
| pCS2-BACH1-GFP-IRES-mCherry | This paper | N/A |
| pCS2-BACH2-GFP-IRES-mCherry | This paper | N/A |
| pCS2-FNIP1degron-GFP-OMP25-IRES-mCherry | 36 | N/A |
| pCS2-COA4-GFP-IRES-mCherry | This paper | N/A |
| pCS2-OXR1-GFP-IRES-mCherry | This paper | N/A |
| pCS2-DHCR7-GFP-IRES-mCherry | This paper | N/A |
| pCS2-CBFB-GFP-IRES-mCherry | This paper | N/A |
| pCS2-PDE3B-GFP-IRES-mCherry | This paper | N/A |
| pCS2-KCTD5-GFP-IRES-mCherry | This paper | N/A |
| pCS2-GFP-TNFAIP3-IRES-mCherry | This paper | N/A |
| pCS2-RAP1GAP-GFP-IRES-mCherry | This paper | N/A |
| pCS2-GFP-RHOT1-IRES-mCherry | This paper | N/A |
| pCS2-CEP128-GFP-IRES-mCherry | This paper | N/A |
| pCS2-GFP-KCTD2-IRES-mCherry | This paper | N/A |
| pCS2-GFP-RHOT2-IRES-mCherry | This paper | N/A |
| pCS2-PASK-GFP-IRES-mCherry | This paper | N/A |
| pCS2-BACH1-GFP-IRES-mCherry (ΔBTB(Δ1-127), 506-684, Δ506-684) | This paper | N/A |
| pCS2-BACH1-GFP-IRES-mCherry (F9A, Y11F, F125D, F128D, C224S, C299S, C435S, C461S, C492S, C646S, C621S/C625S, L617A/L620A, L630A, Q634E, Y641A) | This paper | N/A |
| lentiV2-puro-sgCTRL | 39 | N/A |
| lentiV2-puro-sgGCSH | This paper | N/A |
| lentiV2-puro-sgBCS1L | This paper | N/A |
| lentiV2-puro-sgNARS2 | This paper | N/A |
| lentiV2-puro-sgPTPMT1 | This paper | N/A |
| lentiV2-puro-sgDLD | This paper | N/A |
| lentiV2-puro-sgMRPL20 | This paper | N/A |
| lentiV2-puro-sgALAD-1 | This paper | N/A |
| lentiV2-puro-sgALAD-2 | This paper | N/A |
| lentiV2-puro-sgFBXO22-1 | This paper | N/A |
| lentiV2-puro-sgFBXO22-2 | This paper | N/A |
| lentiV2-puro-sgALAS1 | This paper | N/A |
| lentiV2-puro-sgHMOX2-1 | This paper | N/A |
| lentiV2-puro-sgHMOX2-2 | This paper | N/A |
| lentiV2-puro-sgCOX10 | This paper | N/A |
| lentiV2-puro-sgCOX15 | This paper | N/A |
| lentiV2-puro-sgCOA7 | This paper | N/A |
| lentiV2-puro-sgCMC2 | This paper | N/A |
| lentiV2-puro-sgPET117 | This paper | N/A |
| lentiV2-puro-sgCOX11 | This paper | N/A |
| pLKO.1-puro-shCTRL | This paper | N/A |
| pLKO.1-puro-shFEM1B | This paper | N/A |
| pLKO.1-puro-shBACH1 | This paper | N/A |
| pET28a-BACH1<sup>CT</sup>-6XHis (wt, L617A/L620A, C646S) | This paper | N/A |
| pMAL-6xHis-MBP-TEV-FEM1B (wt, L18A/L25A, K16E, L18A/L25A/K16E) | This paper | N/A |
| pRSFDuet-1 Elongin B, ElonginC<sup>17-112</sup> | 35 | N/A |
| p6xHis-TEV-Rbx1-T7-6xHis-mysB-TEV-StrepII-CUL2 ($\Delta$ 117-134) | 99 | N/A |
| pCS2-dnCUL2 | Laboratory of Michael Rapé | N/A |
| **Software and algorithms** |  |  |
| GraphPad Prism 10 | GraphPad Software Inc. | RRID:SCR_002798 |
| FlowJo 10.8.1 | FlowJo | RRID:SCR_008520 |
| ImageJ version 1.54r | Fiji | RRID:SCR_002285 |
| Spectronaut version 18.0 | Biognosys | https://biognosys.com/software/spectronaut/ |
| ChimeraX version 5.0 | UCSF | RRID:SCR_015872 |
| Phenix version 2.0 | https://www.phenix-online.org/, PMID: 31588918 | RRID:SCR_014224 |
| Coot version 0.9.8.96 | PMID: 20383002 | RRID:SCR_014222 |
| Octet DataAnalysis software v11.1.0.4 | Sartorius | RRID:SCR_023267 |
| **Other** |  |  |
| ANTI-FLAG M2 Affinity Agarose Gel Slurry | Sigma-Aldrich | Cat# A2220 |
| Amylose Resin | New England Biolabs | Cat# E802 |
| Ni-NTA | QIAGEN | Cat# 30210 |
| Dynabeads™ Protein G for Immunoprecipitation | ThermoFisher | Cat# 10004D |
| SF Cell Line 4D-Nucleofector® X Kit S | Lonza | Cat# V4XC-2032 |
| TransIT®-293 Transfection Reagent | Mirus | Cat# MIR 2704 |
| Opti-MEM | ThermoFisher | Cat# 31985-070 |
| Quantifoil® R 1.2/1.3 Holey Carbon Films on Grids | Electron Microscopy Sciences | Cat # Q310CR1.3 |
| Zeba Spin desalting columns, 7K MWCO, 0.5 mL | ThermoFisher | Cat# 89877 |
| HiLoad 16/600 Superdex 200 pg column | Cytiva | Cat# 28989335 |
| HiLoad 16/60 Superdex 75 prep grade column | Cytiva | Cat# 17106801 |
| Superdex 200 Increase 10/300 GL column | Cytiva | Cat# 28990944 |
| Superose 6 10/300 GL column | Cytiva | Cat# 29091596 |
| 3X FLAG peptide | MedChemExpress | Cat# HY-P0319 |
| HiTrap Q HP anion exchange column | Cytiva | Cat# 17115401 |

---

**Figure S1:**
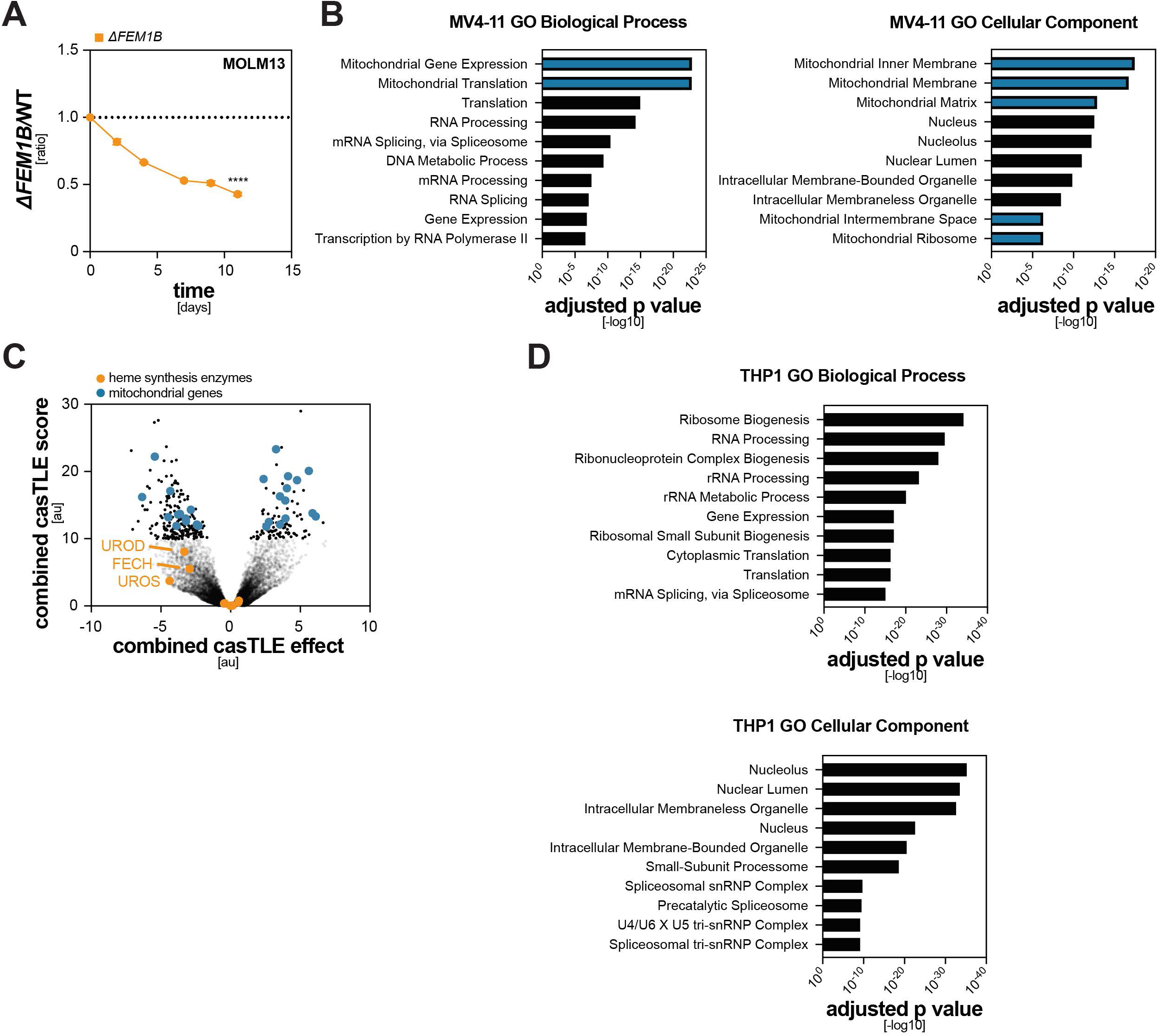
CUL2^FEM1B^ is functionally linked to heme metabolism. Related to Figure 1. **A.** FEM1B promotes AML cell proliferation. mCherry-expressing *ΔFEM1B* MOLM13 cells were mixed with GFP-expressing WT cells and followed by flow cytometry for 12d. Datapoints represent mean ± S.E.M. of n=3 independent experiments. **B.** Mitochondrial processes and components are essential in the MV4-11 cell line used in the synthetic lethality CRISPR screen. Essential genes in MV4-11 WT cells were determined using combined CasTLE analysis of differential sgRNA abundance at d0, d12 and d21 of the whole genome CRISPR screen presented in Figure 1B. Gene list enrichment analysis of <5% FDR scoring depleted genes (n=504 genes) was performed using Enrichr ^88^. **C.** Whole genome dropout CRISPR screen in THP1 cells with low mitochondrial activity does not reveal ties to heme biosynthesis. Combined CasTLE analysis of change in sgRNA abundance over 30 days in two distinct *ΔFEM1B* THP1 clones compared to WT THP1 cells; 10% FDR. **D.** Mitochondrial genes are not enriched among top essential pathways in THP1 cells. Essential genes in THP1 wt cells were determined using a combined CasTLE analysis of differential sgRNA abundance at d0, d21 and d30 of the whole genome CRISPR screen presented in Figure S1C. Gene list enrichment analysis of n=504 highest scoring depleted genes was performed using Enrichr.

**Figure S2:**
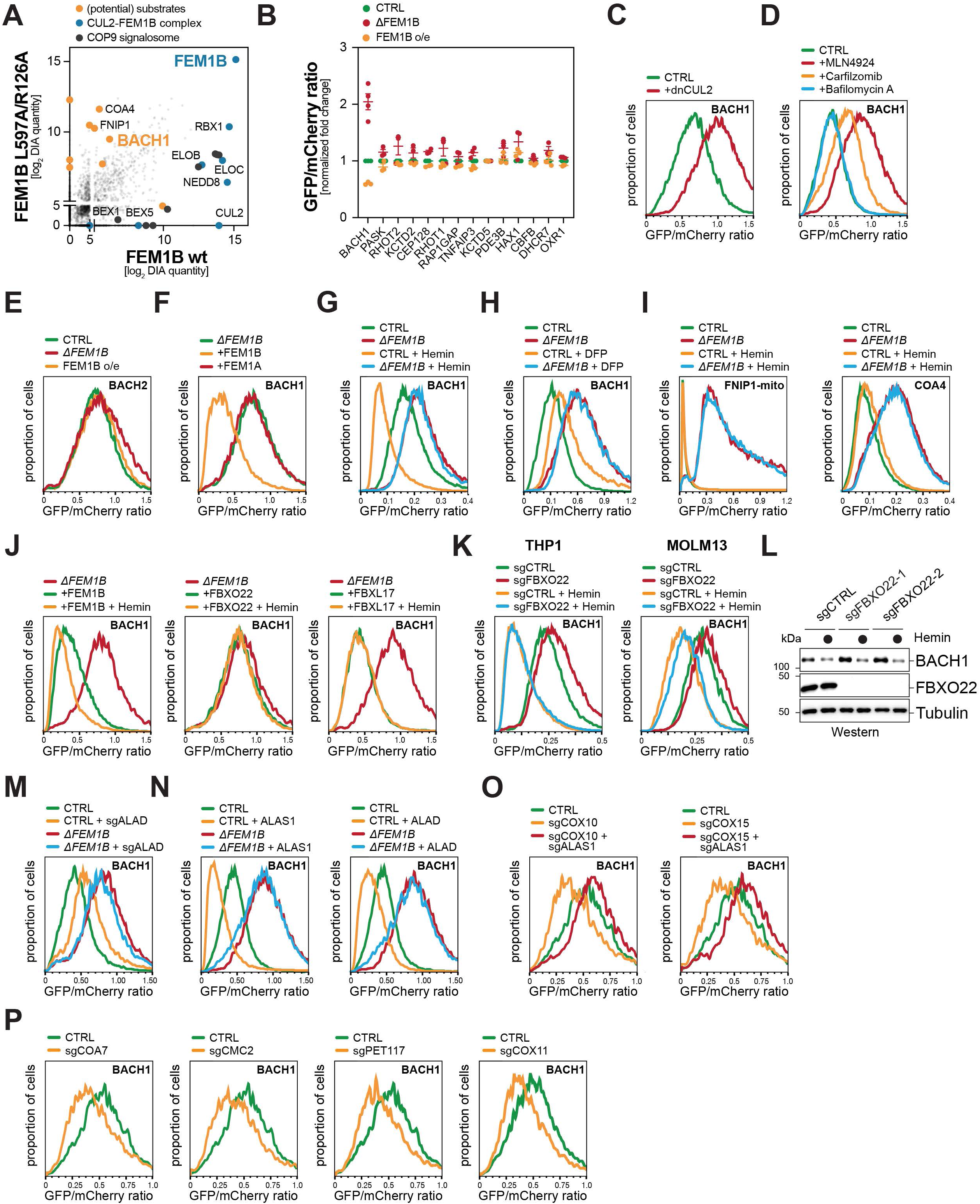
CUL2^FEM1B^ drives heme-dependent degradation of BACH1. Related to Figure 2. **A.** FEM1B binds BACH1. Immunoprecipitation of ^3XFLAG^FEM1B or ^3XFLAG^FEM1B^R126A/L597A^ from THP1 cells coupled with mass spectrometry identifies BACH1 as a candidate substrate of CUL2^FEM1B^. **B.** BACH1 degradation depends on FEM1B. Stability reporters for selected FEM1B binders were transiently expressed in WT or *ΔFEM1B* HEK293T cells in the absence or presence of overexpressed FEM1B. The GFP/mCherry ratio was determined using flow cytometry. Data represented as median ± S.E.M. of median fluorescence intensity ratios (MFI) of n=3-4 independent experiments. **C.** Dominant negative CUL2 stabilizes the BACH1 stability reporter in HEK293T cells. Similar results in n=2 independent experiments. **D.** Inhibition of Cullin-Ring ligases or proteasomes, but not lysosomes, stabilizes BACH1. BACH1 stability reporters were expressed in wt HEK293T cells, carfilzomib (2 μM), bafilomycin A (700 nM) were added for 6h, and MLN4924 (1µM) was added for 16h. Similar results in n=2 independent experiments. **E.** CUL2^FEM1B^ does not target BACH2. BACH2 stability reporters were expressed in WT or *ΔFEM1B* HEK293T cells in the absence or presence of FEM1B. Similar results in n=2 independent experiments. **F.** CUL2^FEM1A^ does not target BACH1. BACH1 stability reporters were expressed in *ΔFEM1B* HEK293T cells in the absence or presence of FEM1B or FEM1A. Similar results in n=2 independent experiments. **G.** Degradation of a BACH1 stability reporter induced by hemin depends entirely on FEM1B in AML cells. BACH1 stability reporters composed of brighter BACH1-mNeonGreen-P2A-mScarlet3 were lentivirally expressed in WT and *ΔFEM1B* MV4-11 cells. Cells were treated with 10µM hemin for 16h. Similar results in n=2 independent experiments. **H.** Iron chelation stabilizes BACH1. BACH1 stability reporters were expressed in WT or *ΔFEM1B* HEK293T cells, which were treated with 1mM DFP for 16h as indicated. Similar results in n=2 independent experiments. **I.** Hemin does not induce degradation of other FEM1B substrates. FNIP1 and COA4 stability reporters were expressed in WT or *ΔFEM1B* HEK293T cells, and cells were treated with 10µM hemin for 16h. Similar results in n=2 independent experiments. **J.** FBXO22 and FBXL17 do not drive heme-induced degradation of BACH1. BACH1 stability reporters were expressed in *ΔFEM1B* HEK293T cells together with FEM1B, FBXO22 or FBXL17, and cells were treated with 10µM hemin for 16h, as indicated. Similar results in n=2 independent experiments. **K.** Depletion of FBXO22 or FBXL17 does not inhibit heme-induced degradation of BACH1 in AML cells. FBXO22 or FBXL17 were depleted from THP1 (left) or MOLM13 (right) cells stably expressing a BACH1 stability reporter. Cells were treated with 10µM hemin for 16h. Similar results in n=2 independent experiments. **L.** Heme-induced degradation of endogenous BACH1 is not affected by depletion of FBXO22. HEK293T cells were depleted of FBXO22 by two distinct sgRNAs. Cells were treated with 10µM hemin for 16h, and endogenous BACH1 levels were assessed by Western blotting. Similar results in n=2 independent experiments. **M.** Depletion of the heme biosynthesis enzyme ALAD stabilizes BACH1 dependent on FEM1B. ALAD was depleted from WT or *ΔFEM1B* HEK293T cells stably expressing a BACH1 stability reporter. Similar results in n=3 independent experiments. **N.** Increasing endogenous heme destabilizes BACH1 dependent on FEM1B. BACH1 stability reporters were expressed in HEK293T cells together with vectors for expression of ALAS1 (left) or ALAD (right). Similar results in n=2 independent experiments. **O.** Disruption of heme integration into ETC cIV destabilizes BACH1 dependent on FEM1B and heme levels. COX10 (left) or COX15 (right) were co-depleted with ALAS1 from WT HEK293T cells stably expressing the BACH1 stability reporter. Similar results in n=2 independent experiments. **P.** Disruption of ETC cIV assembly destabilizes BACH1. cIV assembly factors were depleted from WT HEK293T cells stably expressing the BACH1 stability reporter. Similar results in n=2 independent experiments.

**Figure S3:**
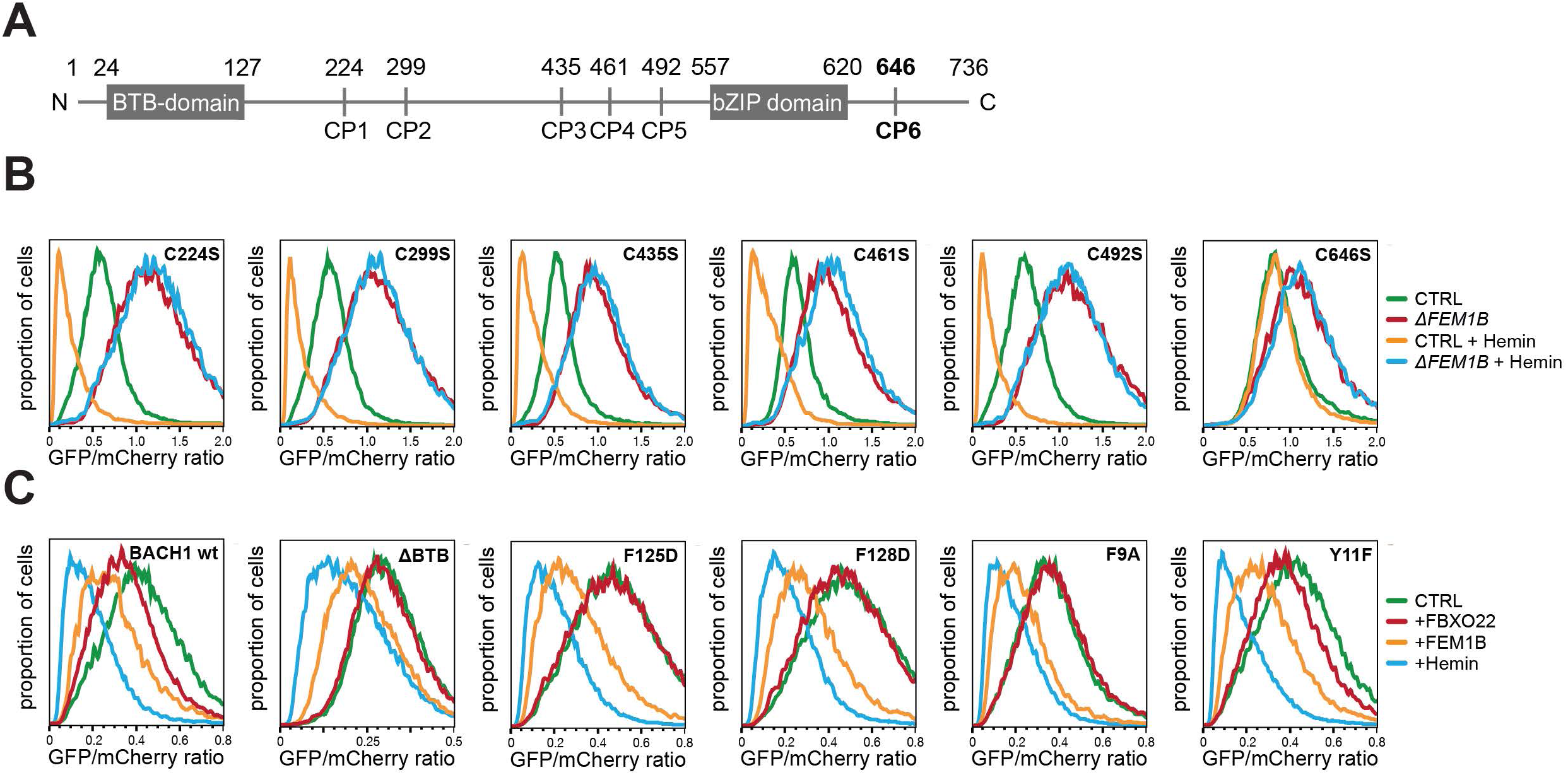
CUL2^FEM1B^ recognizes a specific Cys/Pro motif in BACH1. Related to Figure 3. **A.** Domain map of BACH1 with Cys/Pro motifs (CP) highlighted. **B.** Mutation of the C-terminal CP motif disrupts heme-induced BACH1 degradation through CUL2^FEM1B^. BACH1 stability reporters were expressed in WT and *ΔFEM1B* HEK293T cells. Cells were treated with 10µM hemin for 16h. Similar results in n=2 independent experiments. **C.** Deletion or mutation of the N-terminal BTB-domain does not affect heme-induced BACH1 degradation. BACH1 stability reporters were expressed in WT HEK293T cells together with FEM1B or FBXO22. Cells were treated with 10µM hemin for 16h as indicated. Similar results in n=2 independent experiments.

**Figure S4:**
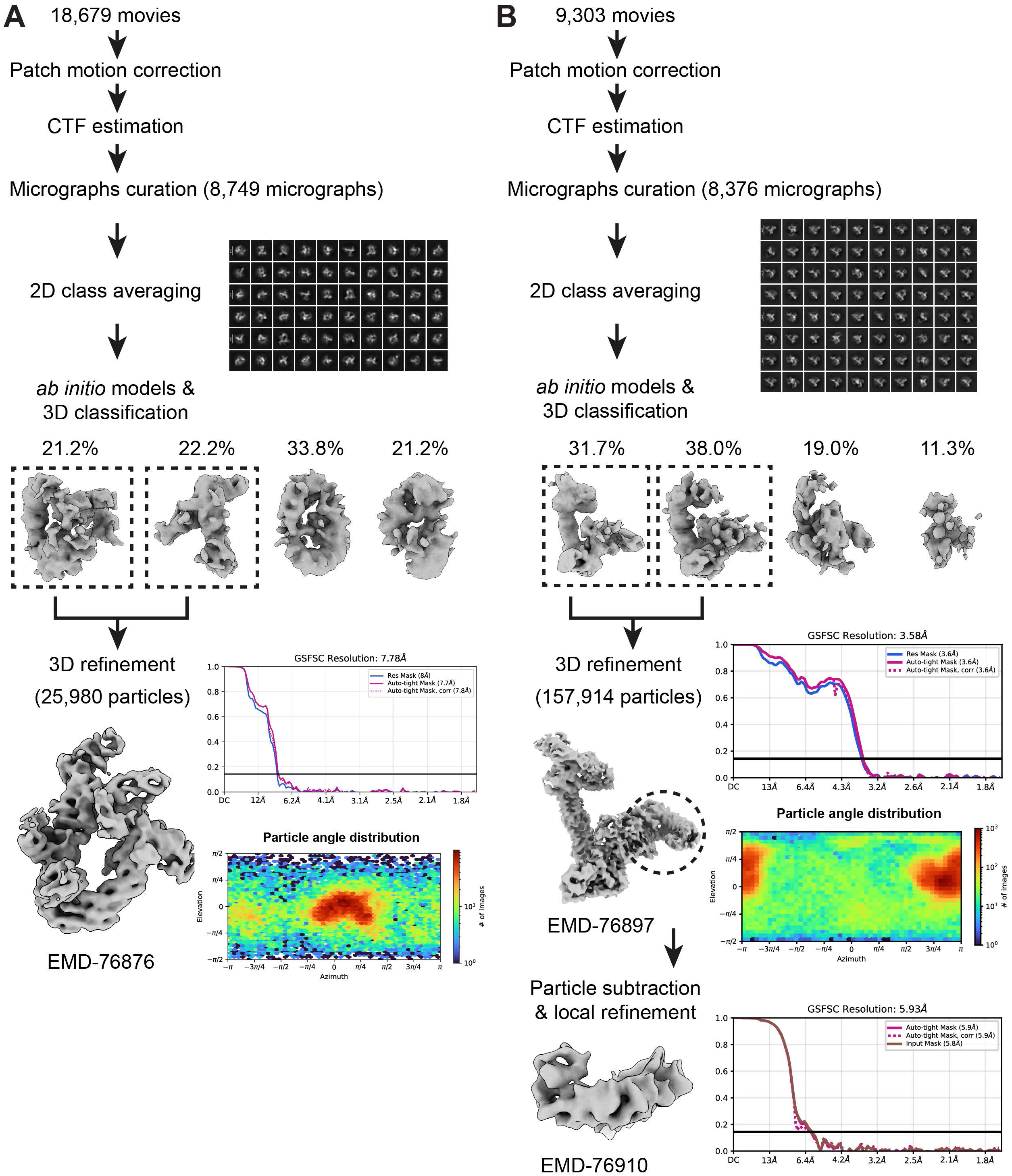
Cryo-EM processing workflow. Related to Figure 3. **A.** Cryo-EM data processing workflow of the CUL2-RBX1-ELOB-ELOC-FEM1B-BACH1^CT^ complex in dimeric conformation (EMD-76876). **B.** Cryo-EM data processing workflow of the CUL2-RBX1-ELOB-ELOC-FEM1B-BACH1^CT^ complex in monomeric conformation (EMD-76897) and local refinement of the FEM1B-BACH1 interface region (EMD-76910) highlighted in dashed line circle in the monomeric conformation map.

**Figure S5:**
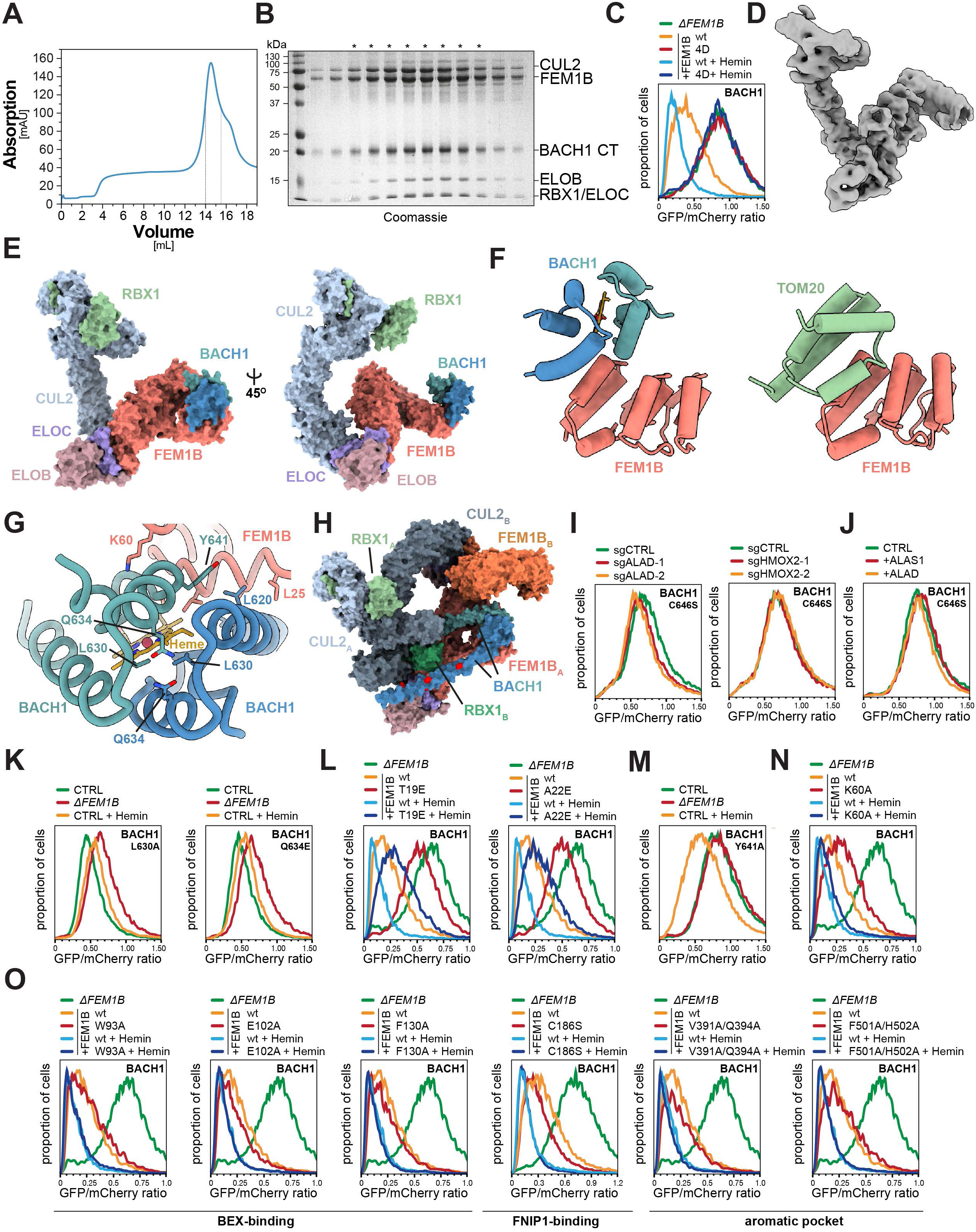
CUL2^FEM1B^ binds BACH1 at its mitochondrial anchor site. Related to Figure 3. **A.** Size exclusion chromatogram of the CUL2-RBX1-ELOB-ELOC-FEM1B-BACH1^CT^ complex. The complex elutes at the volume expected for a dimeric E3 ligase. Collected fractions indicated by a dashed line. Similar results in n>5 independent experiments. **B.** Representative SDS-PAGE analysis of collected fractions reveals a fully assembled complex. (*) denotes fractions collected for structural analysis. Similar results in n>5 independent experiments. **C.** Mutation of FEM1B residues involved in homodimerization (4D) disrupt its ability to induce BACH1 degradation. FEM1B WT or 4D-mutant (F549D/V584D/I587D/L588D) and BACH1 stability reporters were expressed in *ΔFEM1B* HEK293T cells, and cells were treated with 10µM hemin for 16h. Similar results in n=2 independent experiments. **D.** BACH1-bound CUL2^FEM1B^ complex frozen on carbon coated EM-grids yields a monomeric structure. Cryo-EM density map of the CUL2-RBX1-ELOB/C-FEM1B-BACH1^CT^ complex, EMD-76897, contour level 0.052) **E.** Surface representation of the complex shown in (D.) reveals structural details of complex assembly and highlights positioning of BACH1 at the flexible N-terminus of FEM1B (PDB 12ZS). Same colors used as in Figure 3E. Right: 45° rotation to the right. **F.** BACH1 and TOM20 both bind the cap of the FEM1B N-terminal ankyrin repeats. Left: Cryo-EM guided AF3 model of the FEM1B-BACH1 interface. Right: FEM1B-TOM20 interface from the published locally refined structure (PDB 9JCE) ^61^. **G.** Dimerization of BACH1 allows additional interaction with the backside of the FEM1B N-terminus. FEM1B-BACH1 interface highlighting residues L630/Q634 involved in BACH1 dimerization and residues mediating interactions at the backside of FEM1B (BACH1 Y641 and FEM1B K60). **H.** Modeling the extension of BACH1 helices places the lysine rich DNA-binding domains close to RBX1 *in trans*. BACH1 lysine residues in proximity to RBX1 are shown in red. **I.** Depletion of endogenous heme biosynthetic enzymes does not affect BACH1 stability if heme binding has been obliterated. ALAD, or HMOX2 were depleted from WT HEK293T cells stably expressing a BACH1^C646S^ stability reporter. Similar results in n=2 independent experiments. **J.** Increasing endogenous heme does not affect mutant BACH1 degradation. BACH1^C646S^ stability reporters were expressed in HEK293T cells together with ALAS1 or ALAD, as indicated. Similar results in n=2 independent experiments. **K.** Mutation of residues involved in dimerization of BACH1^CT^ renders BACH1 insensitive to FEM1B-mediated and heme induced degradation. BACH1^L630A^ or BACH1^Q634E^ stability reporters were expressed in WT or *ΔFEM1B* HEK293T cells, and cells were treated with 10µM hemin for 16h. Similar results in n=2 independent experiments. Stability of BACH1^wt^ under these conditions is shown in Figure 3K. **L.** Mutation of FEM1B residues at the N-terminal helix disrupt BACH1 degradation. FEM1B variants and BACH1 stability reporters were expressed in *ΔFEM1B* HEK293T cells and cells were treated with 10µM hemin for 16h. Similar results in n=3 independent experiments. **M-N.** Mutation of residues at the backside FEM1B surface recruiting the second BACH1^CT^ dimer subunit attenuate heme-induced degradation of BACH1. WT-FEM1B or FEM1B^K60A^ variants and/or WT-BACH1 or BACH1^Y641A^ stability reporters were expressed in HEK293T cells, which were treated with 10µM hemin for 16h. Similar results in n=2 independent experiments. **O.** FEM1B residues in the central grove required for binding other substrates or regulators are not required for BACH1 degradation. FEM1B variants and BACH1 stability reporters were expressed in *ΔFEM1B* HEK293T cells, which were treated with 10µM hemin for 16h. Similar results in n=2 independent experiments.

**Figure S6:**
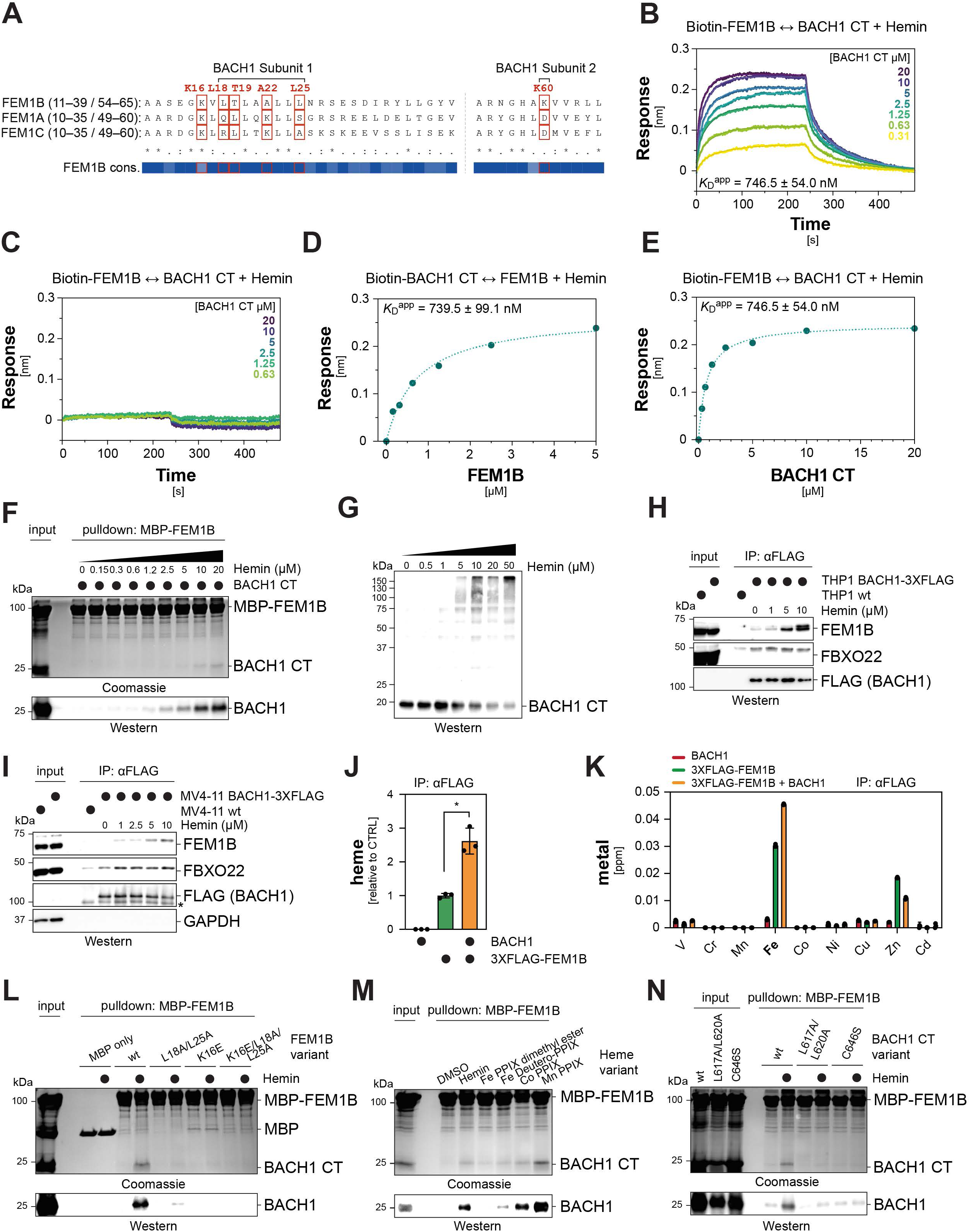
Heme acts as a ternary molecular glue. Related to Figure 4. **A.** FEM1B residues involved in BACH1 binding are highly conserved but specific to FEM1B. Multiple sequence alignment of FEM1A, FEM1B and FEM1C and conservation scores of FEM1B shown for N-terminal FEM1B regions involved in BACH1 binding. Critical residues validated in this study highlighted in red boxes. **B-C.** Binding of FEM1B to BACH1^CT^ requires heme as a molecular glue. Biolayer interferometry using streptavidin tips bound to Biotin-FEM1B were incubated with increasing concentrations of BACH1^CT^ in the presence (B.) or absence (C.) of 20µM Hemin and association/dissociation were monitored over time. Sensorgrams were fit to a 2:1 heterogeneous ligand binding model and fitted curves are overlaid with experimental data. The apparent K_D_ in the presence of 20 µM hemin was determined using steady-state analysis. Similar results in n=2 independent experiments. **D-E.** Apparent dissociation constant for the interaction of Biotin-BACH1^CT^ and FEM1B (D.) or Biotin-FEM1B and BACH1^CT^ (E.) in the presence of 20 µM hemin was determined by plotting the equilibrium response as a function of ligand concentration and fitting to a steady-state binding isotherm. The apparent K_D_ values are reported as best fit ± standard error of the steady-state fit. **F.** Binding of BACH1^CT^ to FEM1B requires heme as a molecular glue. Recombinant BACH1^CT^ was subjected to pulldown assays with MBP-FEM1B immobilized to amylose resin with increasing concentrations of hemin. Interaction was detected by Coomassie staining of SDS-PAGE gels or Western blotting. Similar results in n=2 independent experiments. **G.** Ubiquitylation of BACH1^CT^ by CUL2^FEM1B^ shows a dose-dependent requirement for heme. *In vitro* ubiquitylation assays of recombinant BACH1^CT^ by NEDD8-modified CUL2^FEM1B^ with E1, UBE2R1 and ubiquitin probed for BACH1 ubiquitylation by Western blotting against BACH1^CT^. Similar results in n=2 independent experiments. **H-I.** Interaction of endogenous BACH1 with FEM1B in AML cells depends on heme. BACH1^3XFLAG^ endogenously edited THP1 (H.) or MV4-11 (I.) cells were lysed in the presence of increasing concentrations of hemin and subjected to immunoprecipitation and western blotting with the indicated antibodies. Similar results in n=2 independent experiments. **J.** Purified FEM1B-BACH1 complex contains heme. ^3XFLAG^FEM1B and BACH1^HA^ were transiently expressed in Expi293F cells and lysates subjected to αFLAG-immunoprecipitation, The concentration of heme in eluted protein complexes was determined using an apo-peroxidase based assay. Technical replicates represented as as mean ± SD. Similar results in n=2 independent experiments. Statistical significance was determined using a two-tailed Student’s t-test (* p<0.1). **K.** Purified FEM1B-BACH1 complex contains iron. Eluted proteins obtained as described above were further digested with chymotrypsin and denatured with HNO_3_. Samples were then probed for several transition metals using inductively coupled plasma spectroscopy. Technical replicates represented as mean ± SD. Similar results in n=2 independent experiments. **L.** Mutation of critical FEM1B residues abrogates heme-induced BACH1 binding *in vitro*. Recombinant BACH1^CT^ was subjected to pulldown assays with MBP-FEM1B mutants immobilized to amylose resin ± 20µM hemin. Interaction was detected by Coomassie staining of SDS-PAGE gels or western blotting with a BACH1-antibody. Similar results in n=2 independent experiments. **M.** Loss of FEM1B interacting carboxy groups in heme disrupts BACH1-FEM1B binding *in vitro*. Recombinant BACH1^CT^ was subjected to pulldown assays with MBP-FEM1B mutants immobilized to amylose resin in the presence of 20µM heme or heme analogues. Interaction was detected by Coomassie staining of SDS-PAGE gels or western blotting with a BACH1-antibody. Similar results in n=2 independent experiments. **N.** Mutation of critical BACH1 residues abrogates heme-induced FEM1B binding. BACH1^wt^, BACH1^L617A/L620A^ or BACH1^C646S^ (all BACH1^CT^) were subjected to pulldown assays with MBP-FEM1B immobilized to amylose resin ± 20µM hemin. Interaction was detected by Coomassie staining of SDS-PAGE gels or Western blotting with a BACH1-antibody. Similar results in n=2 independent experiments.

**Figure S7:**
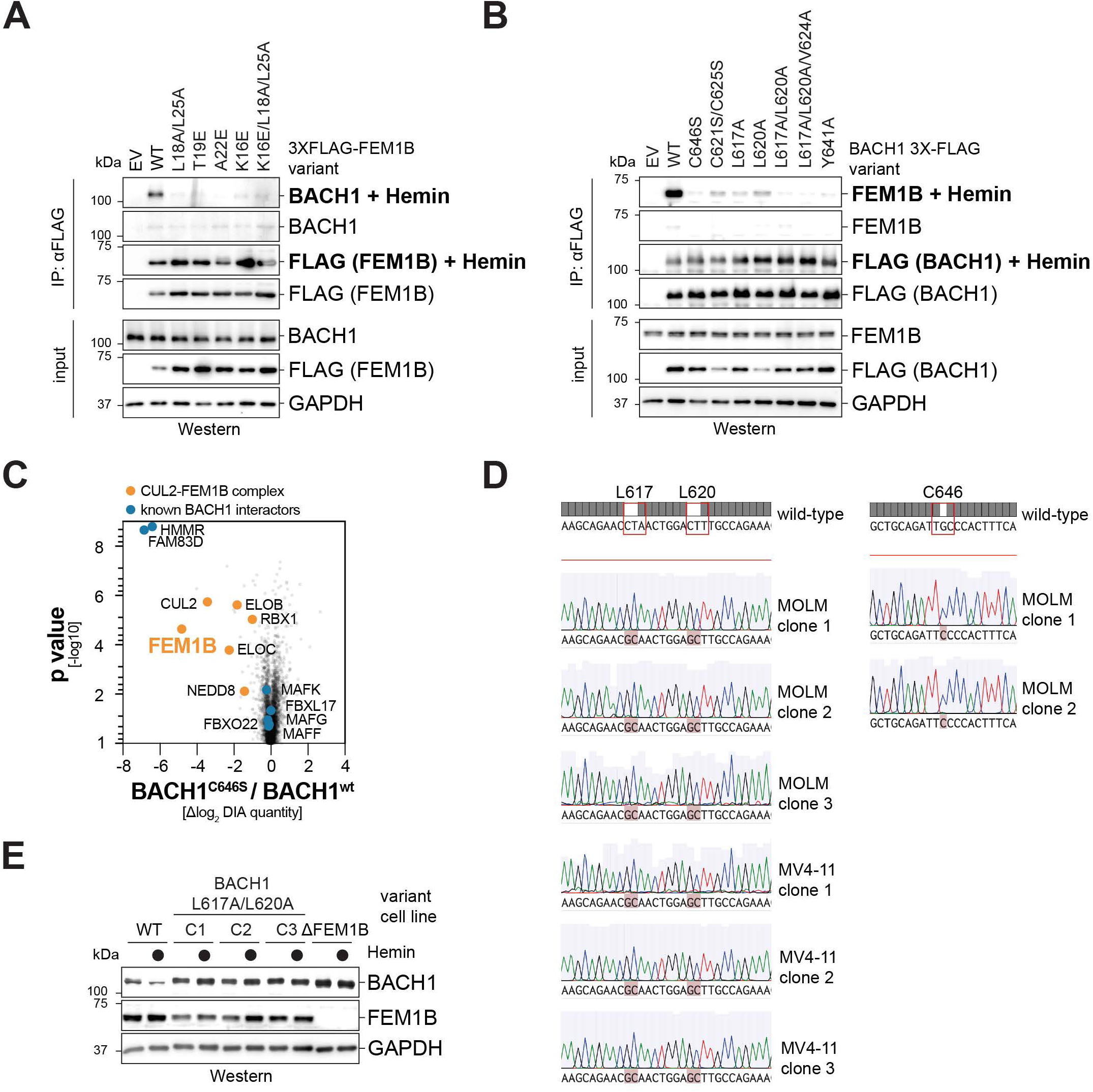
Mutation of critical BACH1 or FEM1B residues in cells disrupts binding and heme-induced degradation. Related to Figure 4. **A-B.** Mutation of BACH1/FEM1B residues abrogates heme-mediated interaction in cells. ^3XFLAG^FEM1B (A.) or BACH1^3XFLAG^ (B.) variants were expressed in HEK293T cells and immunoprecipitates were analyzed by Western blotting with the indicated antibodies. Lysis and IP were performed ± 20µM Hemin as indicated. Similar results in n=2 independent experiments. **C.** Mutation of the heme binding residue C646 in BACH1 disrupts binding to CUL2^FEM1B^, while the interaction with known key interactors, such as MAF proteins, remains intact. BACH^wt^ or BACH1^C646S^ (both 3X-FLAG) were lentivirally expressed in MV4-11 cells and immunoprecipitated proteins were detected by mass spectrometry. Data of n=3 technical MS-replicates shown. **D.** Sanger sequencing confirms correct CRISPR editing at the endogenous BACH1 locus. **E.** Mutation of critical residues at the endogenous BACH1 locus renders it insensitive to heme induced degradation. MV4-11 cells were endogenously edited using CRISPR-Cas9 and homology directed repair, treated with 10µM hemin for 16h, and lysates subjected to Western blot analysis. Similar results in n=2 independent experiments.

**Figure S8:**
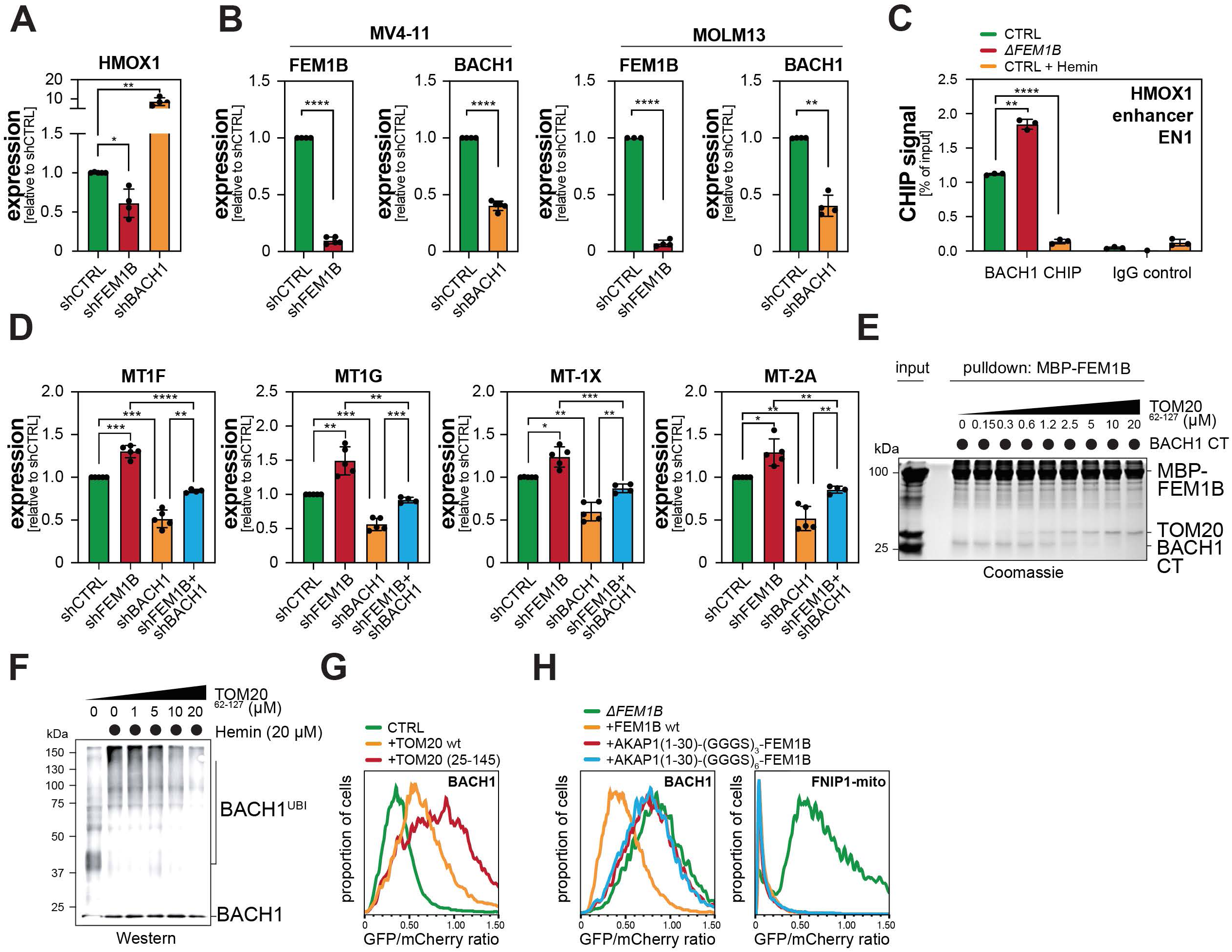
Ternary glue signaling elicits a gene expression response to alleviate heme toxicity. Related to Figure 5. **A.** BACH1 and FEM1B co-regulate HMOX1 expression. BACH1 or FEM1B were depleted in MOLM13 cells using specific shRNAs and HMOX1 expression was analyzed by qPCR. Data of n=4 independent experiments represented as mean ± SD. **B.** FEM1B and BACH1 knockdown verification of MV4-11 and MOLM13 qPCR experiments shown in Figures 5C, 5E and S8A, S8D. Data of n=4-5 independent experiments represented as mean ± SD. **C.** Enhanced BACH1 binding to HMOX1 enhancer upon FEM1B loss. *ΔFEM1B* and wt MV4-11 cells were subjected to BACH1 CHIP and qPCR of known BACH1 binding sites in the EN1 enhancer of HMOX ^89^. Heme treated cells that degrade BACH1 served as control. Data expressed as percentage of input. Similar results observed in n=3 independent experiments. **D.** Metallothionein gene expression is impacted by BACH1- and/or FEM1B-depletion. BACH1 and/or FEM1B were depleted in MV4-11 cells using specific shRNAs and expression of MT1F, MT1G, MT1X or MT2A were monitored by qPCR. Data of n=4-5 independent experiments represented as mean ± SD. **E.** BACH1 and TOM20 compete for binding to FEM1B. Recombinant BACH1^CT^ was subjected to pulldown assays with MBP-FEM1B immobilized to amylose resin in the presence of 20µM hemin and increasing concentrations of the FEM1B-binding cytosolic domain of TOM20 lacking flexible Lys residues (TOM20^62–127^) ^36^. Interaction was detected by Coomassie staining. Similar results in n=2 independent experiments. **F.** Ubiquitylation of BACH1^CT^ by CUL2^FEM1B^ is impeded by TOM20. *In vitro* ubiquitylation assays of BACH1^CT^ by NEDD8-modified CUL2^FEM1B^ complexes with E1, UBE2R1 and ubiquitin and 20µM hemin. Increasing concentrations of TOM20^62–127^ were added as indicated. BACH1 ubiquitylation was detected by Western blotting. Similar results in n=2 independent experiments. **G.** TOM20 overexpression stabilizes BACH1 in cells. BACH1 stability reporters were expressed in HEK293T cells together with TOM20 or cytoplasmic TOM20^25–145^ as indicated. Similar results in n=2 independent experiments. **H.** Anchoring all FEM1B to mitochondria inhibits BACH1 degradation. BACH1 or FNIP1-mito reporter constructs were transiently transfected into *ΔFEM1B* HEK293T cells together with control, FEM1B^wt^ or FEM1B constructs engineered to be anchored to the outer mitochondrial membrane. The N-terminal anchor helix (residues 1-30) of the OMM protein AKAP1 was appended to the FEM1B N-terminus separated by GGGS-linkers of different lengths to induce mitochondrial tethering. Similar results in n=2 independent experiments. Statistical significance in (A-D.) was determined using one-sample t-tests and two-tailed Student’s t-tests (* p<0.1, ** p<0.01, *** p<0.001 and **** p<0.0001).

**Figure S9:**
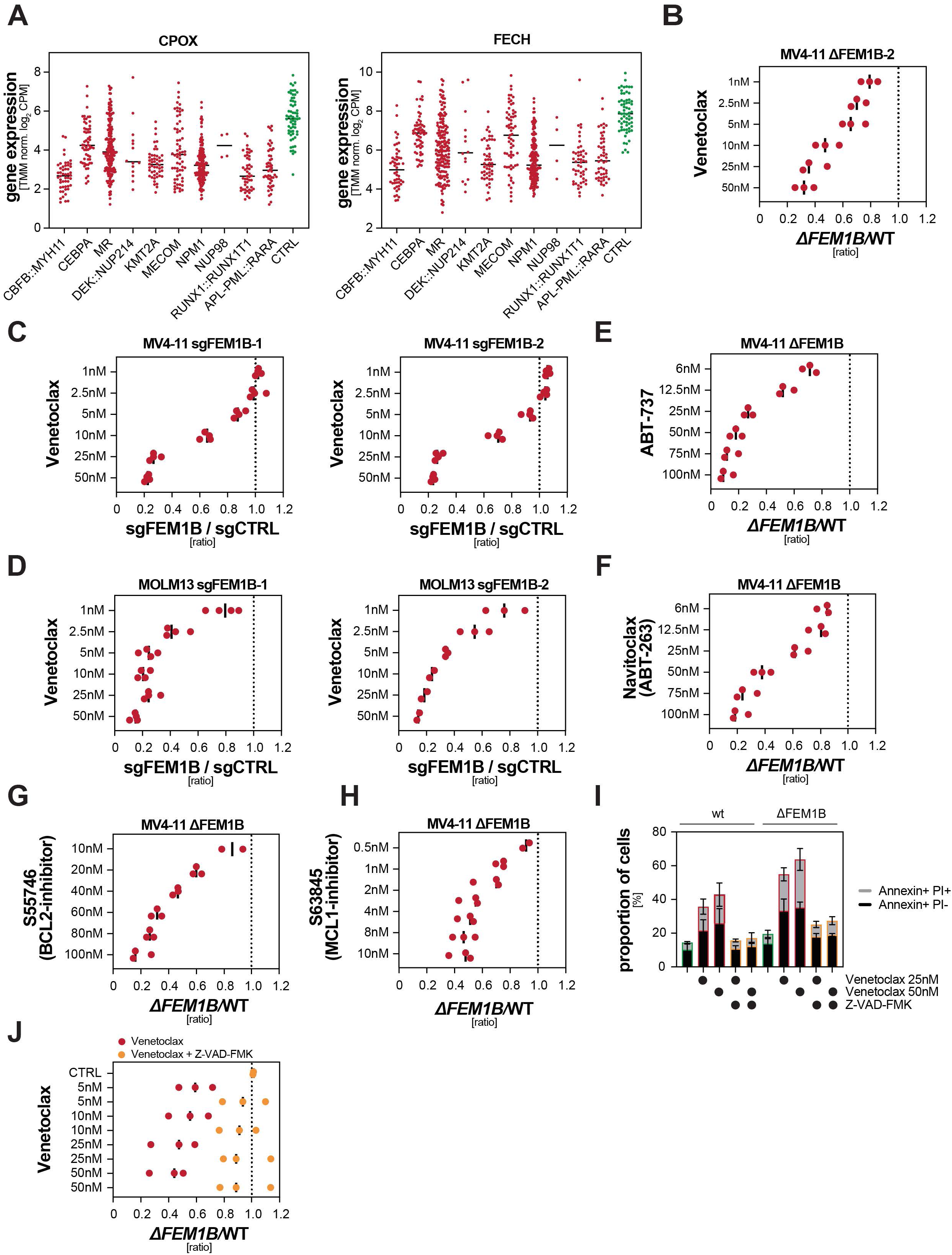
Loss of FEM1B sensitizes AML cells to the BCL2-inhibitor Venetoclax. Related to Figure 6. **A.** Low expression of the representative heme biosynthesis enzymes CPOX and FECH is conserved over all AML subtypes. Gene expression analysis of patients and controls described above separated by genetically defined subtypes according to the WHO 2022 classification. Individual datapoints shown together with mean. **B.** Cell competition assays confirm strong synergism between Venetoclax and loss of FEM1B. mCherry-expressing *ΔFEM1B* and GFP-expressing WT MV4-11 cells were mixed and incubated with the indicated concentrations of Venetoclax for 4 days, after which mCherry/GFP ratios were determined by flow cytometry. Datapoints represent n=3 independent experiments. **C-D.** Acute FEM1B depletion also sensitizes AML cells to Venetoclax treatment. mCherry- or GFP-expressing MV4-11 (C.) or MOLM13 (D.) were infected with sgCTRL or two distinct sgRNAs targeting FEM1B, respectively. Following selection with puromycin, cells were mixed and treated with the indicated concentrations of Venetoclax for 4 days, after which mCherry/GFP ratios were determined by flow cytometry. Datapoints represent n=2-4 independent experiments. **E-H.** Loss of FEM1B sensitizes AML cells to a range of BCL family inhibitors. mCherry-expressing *ΔFEM1B* and GFP-expressing WT MV4-11 cells were mixed and incubated with the indicated concentrations of ABT-737 (E.), Navitoclax/ABT-263 (F.), S55746 (G.) or S63845 (H.) for 4 days, after which mCherry/GFP ratios were determined by flow cytometry. Datapoints represent n=2-3 independent experiments. **I.** *ΔFEM1B* AML cells show increased apoptosis in upon Venetoclax treatment. *ΔFEM1B* and WT MV4-11 cells were treated with the indicated concentrations of Venetoclax for 8 hours and the fraction of early apoptotic (Annexin+/PI-) and late apoptotic (Annexin+/PI+) cells was determined by flow cytometry. Apoptosis was inhibited by addition of 50µM Z-VAD-FMK as indicated. Data expressed as mean ± SD of n=3 independent experiments. **J.** Caspase inhibition rescues synthetic lethality of Venetoclax in *ΔFEM1B* AML cells. mCherry-expressing *ΔFEM1B* and GFP-expressing WT MV4-11 cells were mixed and incubated with the indicated concentrations of Venetoclax and 50µM of Z-VAD-FMK for 2 days, after which mCherry/GFP ratios were determined by flow cytometry. Datapoints represent n=2-3 independent experiments.

